# An ACE2-blocking antibody confers broad neutralization and protection against Omicron and other SARS-CoV-2 variants

**DOI:** 10.1101/2022.02.17.480751

**Authors:** Wenjuan Du, Daniel L. Hurdiss, Dubravka Drabek, Anna Z. Mykytyn, Franziska K. Kaiser, Mariana González-Hernandez, Diego Muñoz-Santos, Mart M. Lamers, Rien van Haperen, Wentao Li, Ieva Drulyte, Chunyan Wang, Isabel Sola, Federico Armando, Georg Beythien, Malgorzata Ciurkiewicz, Wolfgang Baumgärtner, Kate Guilfoyle, Tony Smits, Joline van der Lee, Frank J.M. van Kuppeveld, Geert van Amerongen, Bart L. Haagmans, Luis Enjuanes, Albert D.M.E. Osterhaus, Frank Grosveld, Berend-Jan Bosch

**Author notes:** State Key Laboratory of Agricultural Microbiology, College of Veterinary Medicine, Huazhong Agricultural University, Wuhan, P.R. China.

## Abstract

The ongoing evolution of SARS-CoV-2 has resulted in the emergence of Omicron, which displays striking immune escape potential. Many of its mutations localize to the spike protein ACE2 receptor-binding domain, annulling the neutralizing activity of most therapeutic monoclonal antibodies. Here we describe a receptor-blocking human monoclonal antibody, 87G7, that retains ultrapotent neutralization against SARS-CoV-2 variants including the Alpha, Beta, Gamma, Delta and Omicron (BA.1/BA.2) Variants-of-Concern (VOCs). Structural analysis reveals that 87G7 targets a patch of hydrophobic residues in the ACE2-binding site that are highly conserved in SARS-CoV-2 variants, explaining its broad neutralization capacity. 87G7 protects mice and/or hamsters against challenge with all current SARS-CoV-2 VOCs. Our findings may aid the development of sustainable antibody-based strategies against COVID-19 that are more resilient to SARS-CoV-2 antigenic diversity.

**One sentence summary:** A human monoclonal antibody confers broad neutralization and protection against Omicron and other SARS-CoV-2 variants

## Main Text

Since its emergence in humans late 2019, SARS-CoV-2 has caused >400 million infections and >5.8 millions confirmed deaths worldwide. This massive propagation has allowed rapid evolution of the virus, leading to the independent emergence of a multitude of variants beginning in late 2020. Five of these have been declared by WHO as variants of concern (VOCs) – B.1.1.7 (Alpha), B.1.351 (Beta), P.1 (Gamma), B.1.617.2 (Delta) and B.1.1.529 (Omicron) – as they display increased transmission, immune evasion and/or enhanced disease. Other variants that have spread less widely, but with mutations like those present within VOCs, have been defined as variants of interest (VOIs) such as C.37 (Lambda) and B.1.621 (Mu) (*1*). Some SARS-CoV-2 variants – in particular Beta, Gamma and Omicron – have accrued mutations in the spike (S) protein that correlate with escape from humoral immunity. Sera from patients infected with the ancestral strain and sera from COVID-19 vaccinees exhibit 3 to 9-fold reductions in neutralization activity against Beta and Gamma (*1–3*), whereas neutralizing activity against the globally emerging Omicron was reduced to about 25 to 40-fold (*4–11*). With global population seroprevalence increasing due to natural infection and/or vaccination, the ongoing evolution of SARS-CoV-2 may lead to continuous emergence of antigenically drifted variants that jeopardize the effectiveness of vaccines and antibody-based therapeutics.

Entry of SARS-CoV-2 into host cells is mediated by the trimeric S glycoprotein that consists of two subunits: S1 and S2. The S1 subunit binds the host angiotensin-converting enzyme 2 (ACE2) receptor and the S2 subunit accomplishes membrane fusion. The N-terminal domain (NTD) and the receptor-binding domain (RBD), within the S1 subunit, are the major targets of neutralizing antibodies. These domains are hotspots for mutations observed in SARS-CoV-2 variants that enable escape of serum neutralizing antibodies from infected or vaccinated individuals and of NTD- and RBD-directed monoclonal antibodies. Escape mutations in the RBD are concentrated in the four major and structurally defined neutralizing epitope classes in the RBD(*12*). In particular, the spike proteins of the emerging Omicron BA.1 and BA.2 subvariants carry an unprecedented set of mutations (approximately 30 substitutions, deletions, or insertions) with amino acid substitutions in each of these neutralizing epitope classes, including K417N (class 1), E484A (class 2), G446V (class 3) and G339D (class 4), as well as mutations in the major neutralizing epitope in the NTD (e.g. G142D and deletion of residues 143-145, NTD supersite), potentiating viral escape from vaccine- and infection-elicited antibody-mediated immunity (*13–17*). The escape mutations also have a devastating effect on neutralization by the potent neutralizing ACE2-blocking antibodies corresponding to those that are emergency use authorized for treatment of COVID-19. REGN10933 and REGN10987 (Regeneron), LY-CoV555 and LY-CoV016 (Eli Lilly) completely lost neutralization of Omicron, whereas COV2-2130 and COV2-2196 (parent mAbs of AZD1061 and AZD8895, AstraZeneca) showed an intermediate 12 to 428-fold and 74 to 197-fold loss in neutralization potential against BA.1 Omicron, respectively (*18*). Of the clinically approved or authorized antibodies, S309 (parent of the clinical mAb VIR-7831, Vir Biotechnology) retained significant neutralization against BA.1 Omicron (2 to 3-fold potency loss) but its potency was significantly reduced against BA.2 Omicron (27-fold potency loss) (*15, 16, 18-23*). In general, Omicron escapes existing SARS-CoV-2 neutralizing antibodies with few exceptions, which has major consequences for antibody-based treatment strategies for COVID-19 (*15, 16, 18-22, 24*). Isolation and in-depth characterization of broadly neutralizing and protective antibodies can inform the development of improved vaccines and monoclonal antibody treatments for COVID-19 that are more resistant to antigenically drifted SARS-CoV-2 variants.

Here we identify a SARS-CoV-2 neutralizing human monoclonal antibody, 87G7, with a remarkable broad-spectrum neutralization and protection efficacy. 87G7 blocks SARS-CoV-2 infection via ACE2 binding inhibition with robust neutralizing activity against Alpha, Beta, Gamma, Delta and Omicron. Structural elucidation reveals that 87G7 can bind the highly divergent ACE2 receptor binding site by targeting a patch of hydrophobic residues in convex tip of the receptor-binding ridge that are highly conserved in SARS-CoV-2 variants, including the five VOCs. We demonstrate *in vivo* prophylactic and therapeutic activity by 87G7 against ancestral and variant SARS-CoV-2 using two animal disease models.

### Human monoclonal antibody 87G7 potently neutralizes Omicron and other VOCs/VOIs

To identify human monoclonal antibodies with broad neutralizing capacity against SARS-CoV-2 variants, we explored the antibody repertoire of Harbour H2L2 mice immunized with the SARS-CoV-2 spike protein. The transgenic H2L2 mice encoding chimeric immunoglobulins with human variable heavy and light chains and murine constant region were immunized with plasmid DNA encoding the spike ectodomain and with purified trimeric S ectodomain of the ancestral SARS-CoV-2 strain Wuhan-Hu-1. Hybridoma supernatants with S ectodomain ELISA-reactivity were screened for neutralizing activity against SARS-CoV-2 S pseudovirus with the S E484K mutation, a residue that is at variance in several SARS-CoV-2 VOIs and VOCs playing a key role in resistance to neutralizing antibodies (**Figure 1a**). Among the ∼300 hybridoma supernatants tested, the 87G7 hybridoma supernatant displayed the most potent neutralizing activity. The chimeric 87G7 H2L2 antibody was subsequently converted to a fully human immunoglobulin, by cloning of the human variable heavy and light chain regions into a human IgG1/kappa chain backbone, and the recombinantly expressed 87G7 human monoclonal antibody was evaluated for its capacity to neutralize the prototypic Wuhan-Hu-1 SARS-CoV-2 and the Alpha, Beta, Delta and Omicron VOCs using the VSV pseudovirus neutralization assay. Two therapeutic mAbs REGN10933 and REGN10987 were used for comparison (*25*). 87G7 exhibited potent neutralizing efficacy against Wuhan-Hu-1 S mediated cell entry with a half maximum inhibitory concentration (IC50) of 5.4 ng/ml. In addition, entry of VSV pseudotypes harboring S proteins from VOCs including Alpha, Beta, Delta and Omicron (BA.1 subvariant) was blocked with IC50 values ranging from 1.4 to 5.1 ng/ml. REGN10933 showed decreased neutralization potency against Beta and Omicron corresponding with a 20-fold and 350-fold loss in IC50, respectively, whereas neutralization potential against Omicron by REGN10987 was lost (**Figure 1b** **and 1e)**. Neutralization potency of 87G7 and REGN10933 was subsequently tested in live virus neutralization assay. Relative to an early pandemic strain with D614G spike mutation (D614G), REGN10933 exhibited a fold loss in inhibitory activity against Beta and Gamma of 7.8 and 15.9, respectively, and had fully lost its neutralization potential against Omicron BA.1. In contrast, 87G7 potently neutralized D614G (IC50: 5.7 ng/ml), as well as Alpha, Beta, Gamma, Delta and Omicron (subvariants BA.1 and BA.2) VOCs with IC50 values ranging from 3.1 to 12.5 ng/ml (**Figure 1c** **and 1e)**. In addition, 87G7 neutralized Lambda and Mu variants of interest with similar potency (IC50s: 1.2 and 4.8 ng/ml, respectively) as to D614G (**Figure 1d** **and 1e)**.

**Figure 1.**
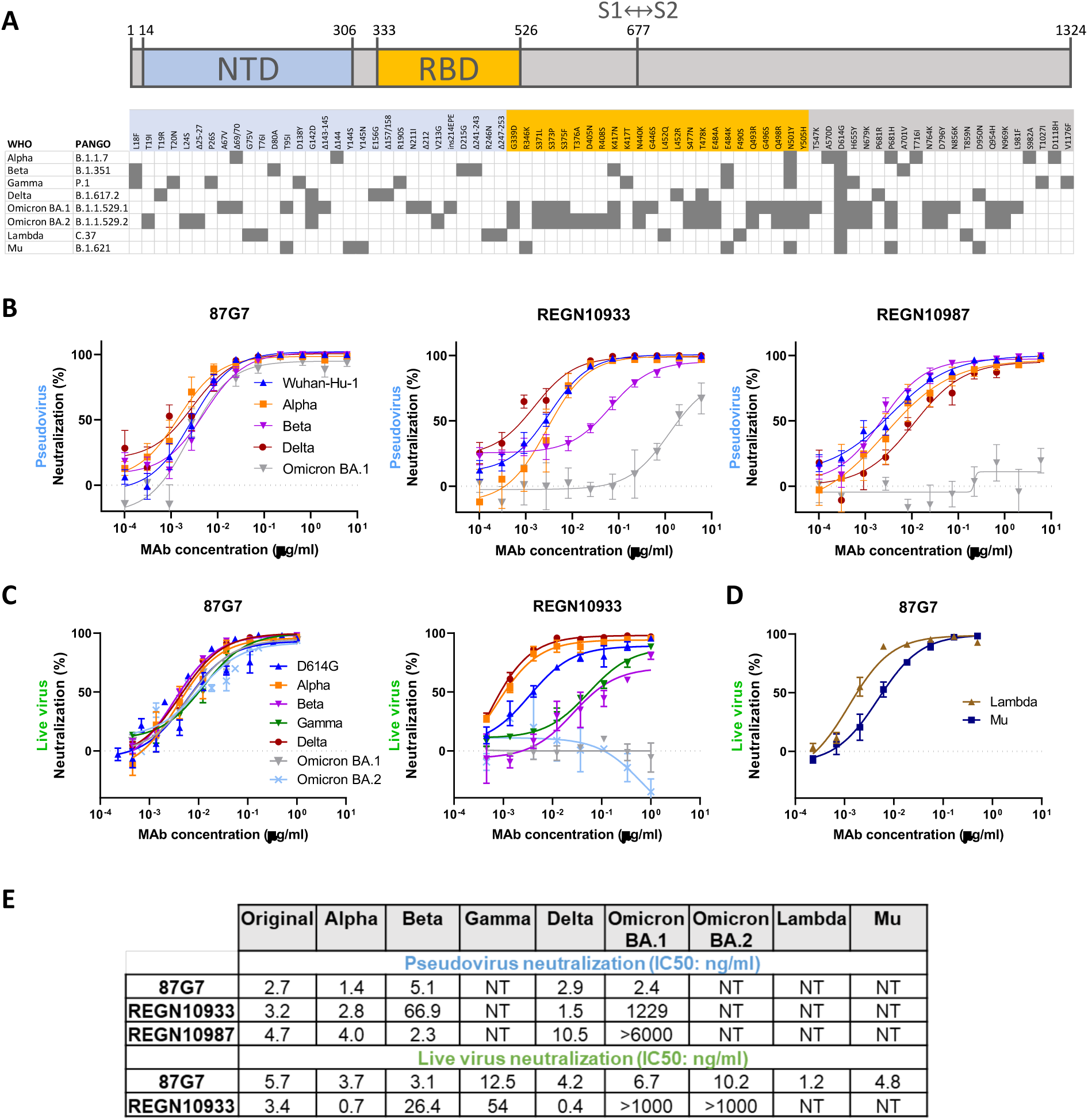
87G7 potently neutralizes Omicron and other SARS-CoV-2 variants. **(A)** S protein schematic with mutations indicated that are found in Alpha, Beta, Gamma, Delta and Omicron (BA.1 and BA.2) variants of concern (VOCs), and Lambda and Mu variants of interest (VOIs), relative to ancestral SARS-CoV-2. The S N-terminal domain (NTD, in blue) and receptor-binding domain (RBD, in orange), and S1/S2 junction are indicated. SARS-CoV-2 lineage naming according to WHO (World Health Organization) and PANGO (Phylogenetic Assignment of Named Global Outbreak). **(B)** Neutralizing activity of 87G7 against virus particles pseudotyped with ancestral SARS-CoV-2 S (Wuhan-Hu-1 strain) or S proteins of Alpha, Beta, Delta and Omicron. Error bars indicate standard deviation between at least two independent replicates. **(C and D)** 87G7 mediated neutralization of live SARS-CoV-2 and variants. Neutralizing potency of 87G7 and REGN10933 against the D614G SARS-CoV-2, and Alpha, Beta, Delta, Gamma and BA.1 and BA.2 Omicron VOCs **(C)** and against Lambda and Mu SARS-CoV-2 VOIs **(D)**. Error bars indicate standard deviation between at least two independent replicates. **(E)** Inhibitory Concentrations 50% (IC50) of 87G7 against SARS-CoV-2 variants calculated from the neutralization curves displayed in panel b, c and d. NT: not tested.

We next evaluated the epitope location and mechanism of action of 87G7. The antibody binds to the S receptor-binding domain (RBD) as demonstrated by ELISA using different SARS-CoV-2 S antigen forms **(Figure S1a)**. By using biolayer interferometry (BLI), we show that 87G7 IgG shows strong, subnanomolar affinity against monomeric S1 and picomolar apparent binding affinity against trimeric S ectodomain, suggesting bivalent binding to the spike trimer **(Figure S1b)**. Binding competition of 87G7 with published monoclonal antibodies targeting distinct RBD epitopes was determined by BLI. Binding interference for 87G7 was only seen with the class 1 antibody REGN10933 indicating an overlapping binding epitope on the RBD **(Figure S1c)**. To understand the mechanism of virus neutralization, we assessed the antibody interference with spike-mediated receptor-binding activity. Similar to REGN10933, 87G7 was found to block the binding of recombinant S trimer to ACE2, as shown by BLI and ELISA-based assay, rationalizing the potent neutralizing activity by 87G7 **(Figure S1d and S1e)**.

### Structural basis for broad neutralization by 87G7

To understand the structural basis for 87G7-mediated neutralization of SARS-CoV-2, we performed cryo-electron microscopy (cryo-EM) analysis on the 6P-stabilized SARS-CoV-2 S trimer (*26*) in complex with the 87G7 Fab fragment (**Figure S2a-d, Table S1**). Three-dimensional (3D) classification of the data revealed that the S ectodomains had all three RBDs in the open conformation with the 87G7 Fab fragment bound to the flexible, convex tip of the receptor-binding ridge (RBR). Subsequent 3D refinement produced a density map with a global resolution of 2.9 Å (**Figure 2a** **and S2e-g)**. Due to the conformational dynamics of the RBD and the flexible nature of the RBR, the epitope-paratope region was poorly resolved. To improve the interpretation of the 87G7 binding site, focused refinement was performed on the Fab-RBD region of the density map, which improved the local resolution sufficiently to resolve the bulky sidechains which make up the majority of the epitope-paratope interface **(Figure S2h)**. Consistent with our BLI data, the 87G7 epitope overlaps with the ACE2 binding site, preventing receptor engagement through steric hindrance (**Figure 2b**). The 87G7 core epitope consists of residues Y421, L455, F456, F486 and Y489, which form a hydrophobic patch on the RBD (**Figure 2c**). 87G7 buries ∼610 Å^2^ of surface area, with light and heavy chains contributing 48% and 52% of total buried surface area (BSA), respectively. The interaction between 87G7 and the RBR is primarily mediated by the CDR H2-3 and CDR L1 and L3 loops, forming a hydrophobic interface. Of note, RBD residues F486 and Y489 insert into a hydrophobic cleft formed by the sidechains of CDR H2-3 residues Y59 and Y103, Y104 and CDR L3 residues F92 and W94. This interaction is reminiscent of the RBD-ACE2 interaction, where F486 penetrates a deep hydrophobic pocket formed by receptor residues F28, L79, Y83, and L97. The 87G7-RBD interface also appears to be stabilized by several hydrogen bonds. Specifically, the backbone carbonyl groups of RBD residues L455 and G485 interact with H30 (CDR L1) and Y59 (CDR H2), respectively. The sidechain of W94 (CDR L3) is also situated in a manner that it may form a hydrogen bond with Y489 and the backbone carbonyl of F486 (**Figure 2d**). Additional residues outside of the core epitope-paratope interface may also contribute to the interaction between 87G7 and the RBD but were not interpreted further due to limited resolution in these areas. To verify the 87G7 epitope, we evaluated the relative contribution of predicted contact residues on antibody binding. Consistent with the structural data, the F486A mutation strongly reduced 87G7 S binding activity in ELISA (**Figure 2e**). F486A also blocked binding by REGN10933 whereas it had no effect on REGN10987 binding, which is consistent with their reported epitopes (*25*). Alanine substitution of Y489 prevented binding by all three antibodies. F456A mutation only slightly impaired binding by 87G7, whereas a stronger reduction in binding was observed for REGN10933 and REGN10987. We assessed the impact of these alanine substitutes in neutralization escape using the pseudovirus system. Pseudovirus production for S^F456A^ and S^Y489A^ however did not yield infectious virus, consistent with the reported unfavorable impact of these alanine substitutions on ACE2 binding (*27*). The S^F486A^ mutant pseudovirus was inefficiently neutralized by 87G7 (and by REGN10933) (**Figure 2f**), mirroring the ELISA binding data and confirming that F486 is a key residue for 87G7 binding and neutralization.

**Figure 2.**
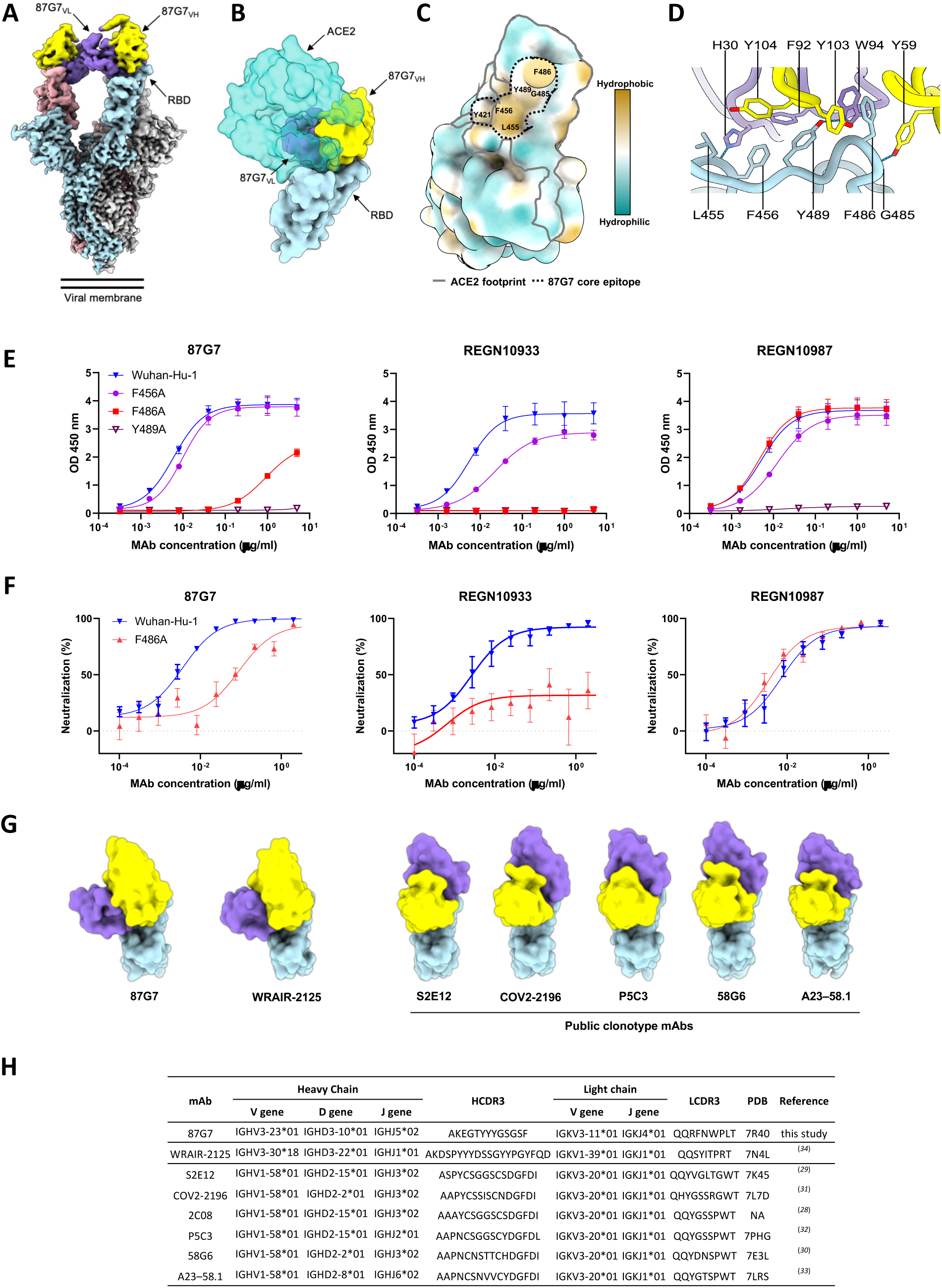
Structural basis for binding and neutralization by 87G7. **(A)** Composite cryo-EM density map for the SARS-CoV-2 spike ectodomain in complex with the 87G7 antibody Fab fragment. The spike protomers are colored blue, gray, and pink, and the 87G7 light- and heavy-chain variable domains colored purple and yellow, respectively. **(B)** Surface representation of the 87G7-bound RBD overlaid with the RBD-bound ACE2 (PDB ID: 6M0J). **(C)** Surface representation of the RBD colored according to the Kyte-Doolittle scale, where the most hydrophobic residues are colored tan and the most hydrophilic residues are colored blue. The residues which make up the 87G7 core epitope and the ACE2 footprint are outlined. **(D)** Close-up view showing selected interactions formed between 87G7 and the SARS-CoV-2 RBD **(E)** ELISA binding of 87G7 to plate-immobilized WT, F456A, F486A and Y489A S1 domains. **(F)** 87G7 neutralizing activity against pseudoviruses with Wuhan-Hu-1 S and S^F486A^. REGN10933 and REGN10987 were taken along as a reference in panel E and F. **(G)** Side-by-side comparison of the SARS-CoV-2 RBD bound to 87G7, WRAIR-2125 (PDB ID: 7N4L), 58G6 (PDB ID: 7E3L), P5C3 (PDB ID: 7PHG), COV2-2196 (PDB ID: 7L7D), S2E12 (PDB ID: 7K45) and A23-58.1 (PDB ID: 7LRS). **(H)** Germline origins of 87G7 and other F486-directed SARS-CoV-2 mAbs with broad neutralization capacity. NA: not applicable.

We next made a structural comparison with other broadly neutralizing mAbs that bind the RBD epitopes with F486 as a key central residue (2C08, 58G6, COV2-2196, P5C3, S2E12 and A23-58.1) (*28–33*). The orientations of these molecules are highly similar, with each binding parallel to the longest axis of the ACE2 binding site and the antibody light chain sitting atop the convex tip of the RBR (**Figure 2g**). These antibodies were derived from different donors but share immunoglobulin heavy (IGHV1-58) and light (IGKV3-20) chain germline origins and display high sequence identity (**Figure 2h**), indicating a public B-cell clone (public clonotype) (*28*). These six public clonotype antibodies potently neutralized Alpha, Beta, Gamma and Delta. Two of them - S2E12 and COV2-2196 - have been assessed thus far for Omicron neutralization, and exhibited a loss in IC50 of 242 (in pseudovirus assay) and 74 to 197-fold (in live virus assay), respectively (*15, 16, 19, 20, 22*). Although 87G7 binds an overlapping epitope and is functionally similar, it has distinct structural and genetic features. Firstly, 87G7 binds perpendicular to the RBR and is rotated ∼122 degrees relative to these other antibodies. Secondly, the ancestral germlines are different and the heavy and light chains have the IGHV3-23 and IGKV3-11 germline origins, respectively. Recently another F486-targeting antibody WRAIR-2125 with broad neutralization potential against Alpha, Beta, Gamma and Delta encoded from distinct heavy-chain (IGHV3-30*18) and light-chain (IGKV1–39*01) germline genes has been reported (*34*). Despite originating from different germline genes, the binding mode is similar for WRAIR-2125 and 87G7, with the aligned Fab:RBD complexes deviating by a root mean square deviation (RMSD) value of 1.9 Å across 200 Cα atom pairs (**Figure 2e**). However, there are differences in the epitope-paratope interactions between these two antibodies. For example, the CDR H3 loop of WRAIR-2125, which is partially disordered in the FAB-RBD crystal structure, is orientated away from the RBR. In contrast, the shorter CDR H3 loop of 87G7 adopts a conformation which places Y103 between F486 and Y489. In addition, the CDR L3 loop of WRAIR-2125 interacts with F486 via T94, whereas the equivalent residue in 87G7 is a tryptophan, creating the possibility for aromatic stacking interactions. Collectively, the sidechains of 87G7 residues Y59 (CDR H2), Y103 (CDR H3) and W94 (CDR L3) create a deep, F486-binding pocket which is not present in WRAIR-2125 **(Figure S3a-b)**. Thus far, WRAIR-2125 has not been assessed for Omicron neutralization.

The 87G7 core epitope residues Y421, L455, F456, F486 and Y489 are highly conserved among SARS-CoV-2 variants (**Figure 3a**). Mutations at these residue positions occur at a very low frequency (< 0.05%) of human-derived SARS-CoV-2 sequences on GISAID as of 5 February 2022. The ACE2 interaction site however comprises a significant number of residues that are mutated in SARS-CoV-2 variants, of which some including K417N, L452R, S477N, T478K, E484A, E484K, F490S, Q493R are close to the 87G7 core epitope and may increase ACE2 affinity and/or enable antibody escape (*25, 27, 35*). We measured the neutralization potential of 87G7 against pseudoviruses carrying S proteins with single site RBD mutations found in VOCs/VOIs. In contrast to REGN10933, 87G7 displayed potent neutralization against all S mutations tested, which is consistent with the ability of 87G7 to retain potent neutralization against the SARS-CoV-2 variants (**Figure 3b**) (*25*).

**Figure 3.**
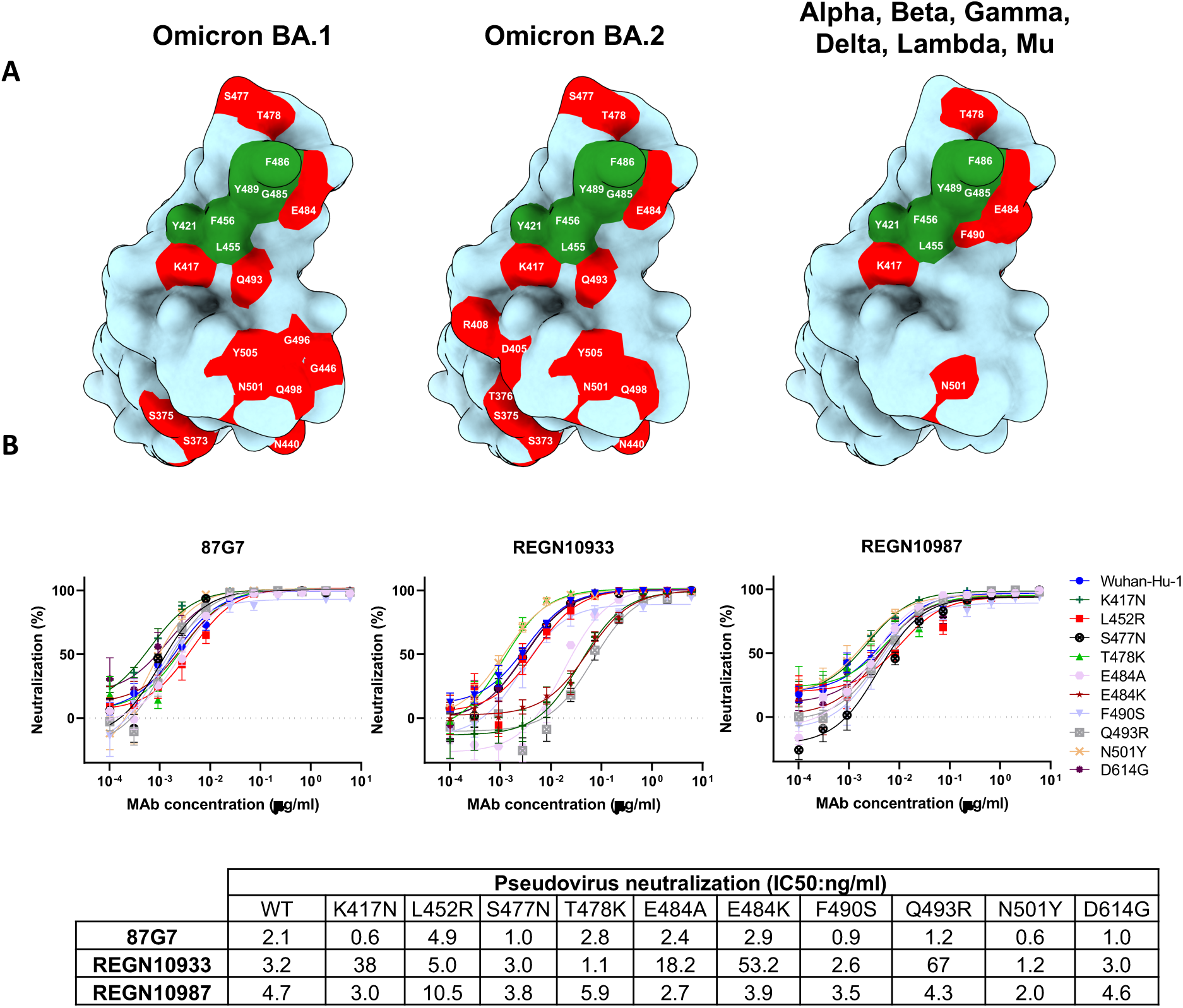
87G7 recognizes a conserved epitope in SARS-CoV-2 RBD. **(A)** Surface representation of the SARS-CoV-2 S RBD with mutations colored red that are found in Omicron BA.1 (left panel) and Omicron BA.2 (middle panel). The right panel displays the set of mutations surrounding the 87G7 core epitope that are present in Alpha, Beta, Gamma, Delta, Lambda or Mu (see also Fig. 1a). The 87G7 core epitope residues are colored green. **(B)** 87G7 neutralizing activity against pseudoviruses with S variants carrying single residue substitutions found in the SARS-CoV-2 variants of concern. The REGN10933 and REGN10987 therapeutic mAbs were used for benchmarking. Data are shown as mean (± SEM) of two independent experiments with technical triplicates, and corresponding IC50 titers are presented in the lower panel.

### 87G7 provides *in vivo* protection from challenge with D614G and SARS-CoV-2 variants

The *in vivo* protection capacity of 87G7 against SARS-CoV-2 challenge was first evaluated using the K18-hACE2 transgenic mice model. To assess prophylactic activity, mice were intraperitoneally injected with 87G7 (10 mg/kg body weight) or an IgG1 isotype control (10 mg/kg) and challenged intranasally 16 hours later with 10^5^ PFU of SARS-CoV-2 using the D614G strain and Alpha, Beta, Gamma or Delta VOCs. To assess therapeutic activity, 87G7 or isotype control was administered (10 mg/kg) at day 1 after challenge with D614G. Mice were scored for weight loss and lungs were collected at day five after challenge for quantification of lung antigen levels and infectious virus. Animals in isotype-treated groups started losing weight after two days (Alpha and Gamma), three days (D614G and Beta) or four days (Delta) post-infection. 87G7-treated animals however were protected from weight loss upon challenge, consistent with the observed reduction in lung antigen levels at day five after challenge in these mice compared to isotype-control treated animals (**Figure 4a** **and b)**. The amount of live virus detected in the lung homogenates decreased by at least one to three orders of magnitude compared to mice receiving the control antibody (**Figure 4c**). Delivery of 87G7 one day after challenge with ancestral virus reduced weight loss (13% of their starting weight relative to 22% in the control group), lung antigen levels, and infectious SARS-CoV-2 titers in lungs by two orders of magnitude (**Figure 4d-f**). These data highlight the prophylactic and therapeutic efficacy in mice by 87G7 against challenge with SARS-CoV-2 and four variants of concern.

**Figure 4.**
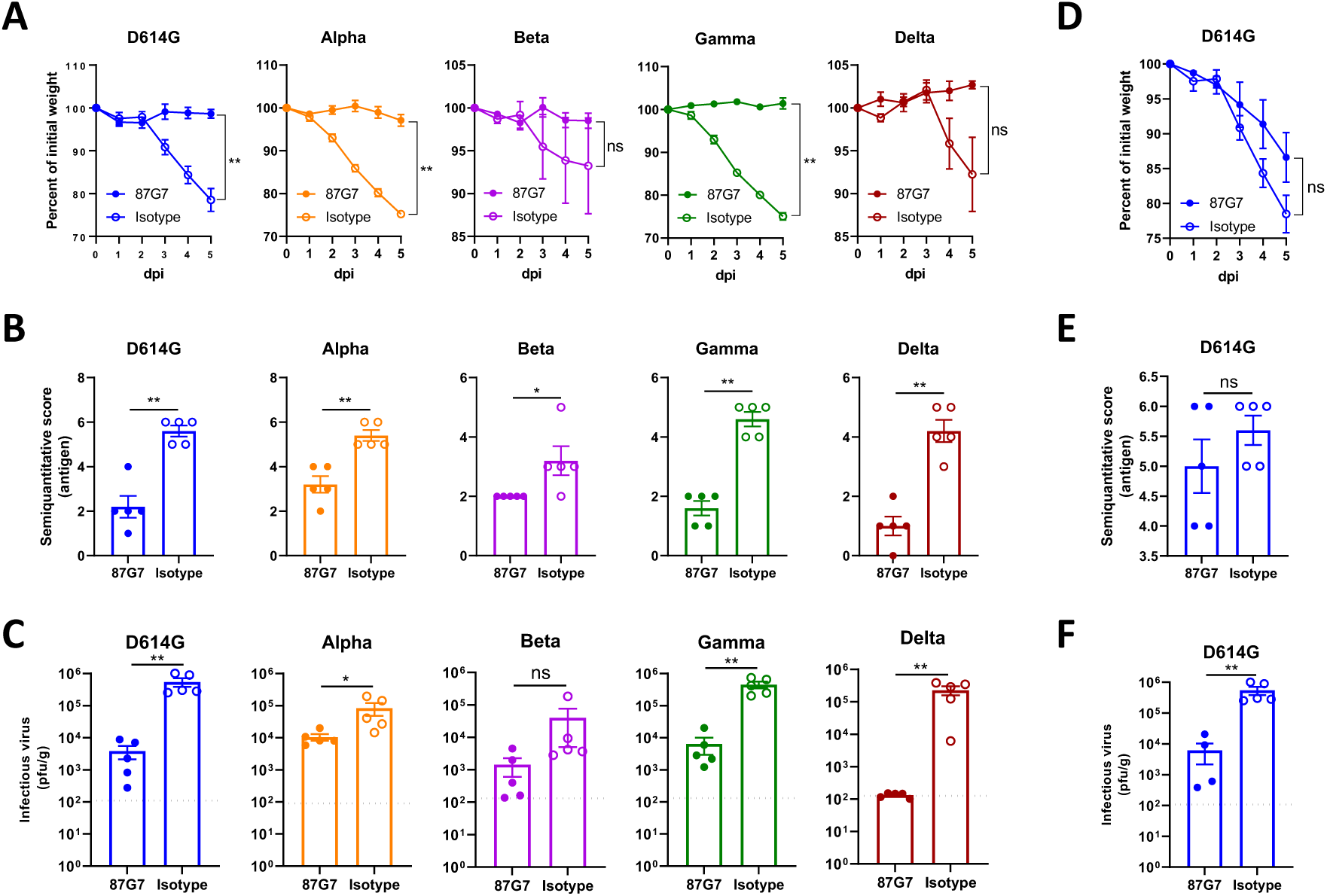
87G7 protects mice from challenge with D614G SARS-CoV-2 and Alpha, Beta, Gamma or Delta variants. Prophylactic and therapeutic treatment was assessed in the K18-hACE2 SARS-CoV-2 mouse model. 87G7 or isotype control mAb was administered intraperitoneally (10 mg/kg body weight) into groups of mice (n = 5) at 24 h before **(A, B, C)** or after virus challenge **(D, E, F)**. Mice were challenged intranasally with 10^5^ PFU of SARS-CoV-2 (D614G, Alpha, Beta, Gamma or Delta) and monitored daily for weight loss (A and D). Five days after challenge lungs were collected from all mice, and lung viral antigen levels were determined by immunohistochemistry **(B and E; Table S2)**, and infectious SARS-CoV-2 loads in lung tissue were measured by plaque assay **(C and G)**. The mean values ± SEM of all data points were shown. Dashed line indicates assay limits of detection. Non-parametric Mann-Whitney U tests were used to evaluate the statistical difference between the 87G7 and isotype-treated groups (**p<0.01, *p<0.05, ns p>0.05).

Protective efficacy by 87G7 was further evaluated in a hamster model of SARS-CoV-2 infection. Syrian hamsters were administered intraperitoneally with 87G7 (10 mg/kg or 20 mg/kg for Omicron-challenged hamsters) or an IgG1 isotype control (10 mg/kg or 20 mg/kg for Omicron-challenged hamsters), 24 h before or 12 h after intranasal challenge with 10^4^ TCID50 of the D614G SARS-CoV-2, Gamma, Delta or Omicron variant. 87G7 administration reduced infectious virus titers in the lungs of most animals in all groups to almost undetectable levels (**Figure 5a**). Preventive treatment with 87G7 reduced infectious virus titers in the nasal cavity of the D614G-, Gamma- and Delta- and Omicron-challenged hamsters by approximately 1-2 logs compared to isotype control antibody treated groups. Histopathological analysis of lung sections from 87G7-treated hamsters showed a markedly reduced number of lesions for all tested variants compared to isotype-treated animals, whereas this pathological difference in the nasal cavity was less prominent (**Figure 5b**). In addition, prophylactic treatment with 87G7 clearly resulted in a notable reduction in antigen expression levels both in the lung and nasal cavity (**Figure 5c**). Therapeutic treatment with 87G7 of the D614G-challenged hamsters significantly reduced infectious virus titers detected in the lungs (>3 log reduction) and nose (approximately 1 log reduction) at day 4 after challenge, and lowered lesions and antigen levels in these respiratory tissues (**Figure 5d-f**).

**Figure 5.**
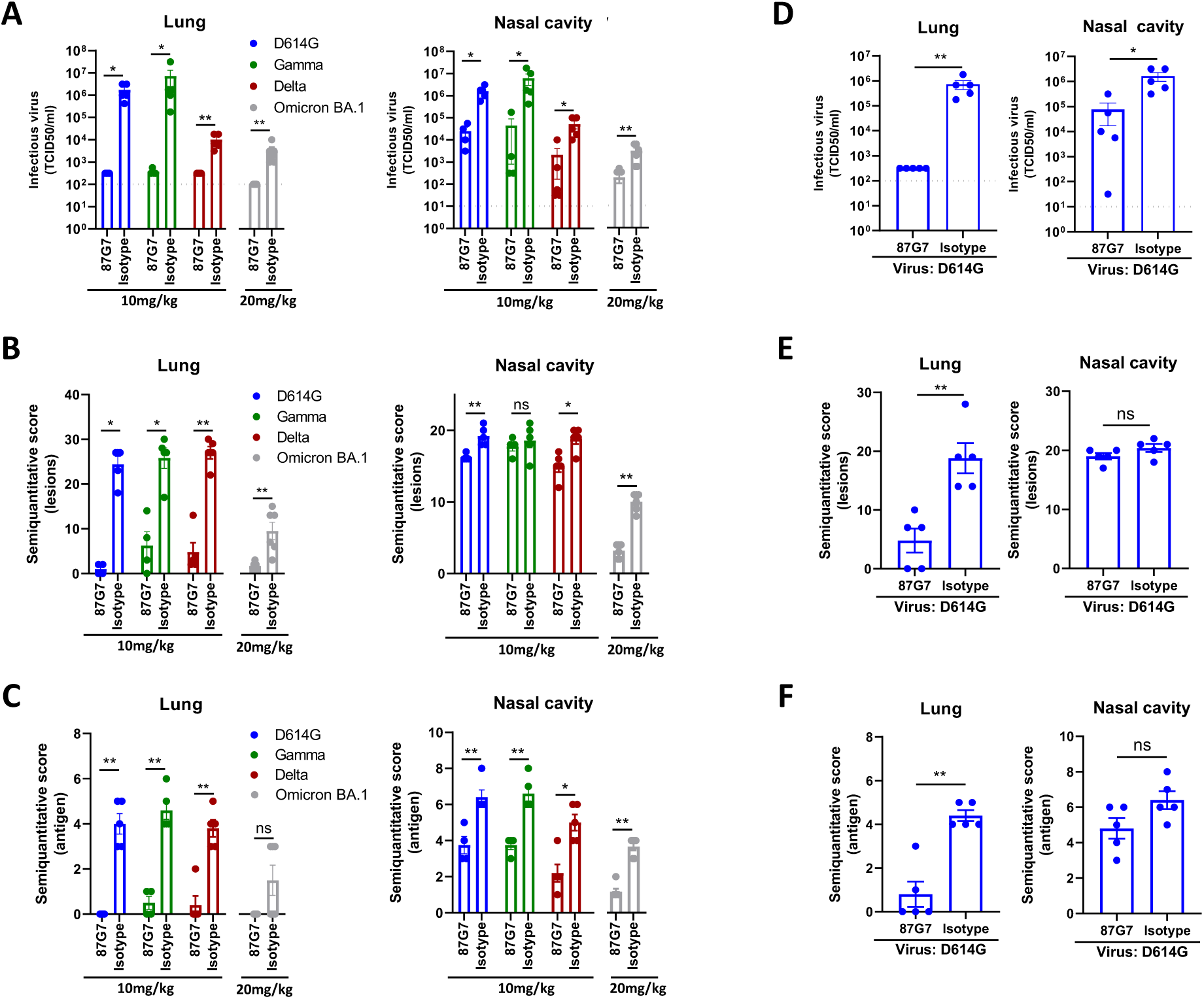
87G7 protects hamsters from challenge with D614G SARS-CoV-2 and Gamma, Delta and Omicron variants. 87G7 or isotype control mAb was administered intraperitoneally (10 mg/kg body weight or 20mg/kg for Omicron-challenged hamsters) into groups of Syrian hamsters (n = 5, and n = 6 for Omicron groups) at 24 h before **(A, B, C)** or 12 h after virus challenge **(D, E, F)**. Hamsters were challenged intranasally with 10^4^ TCID50 of D614G SARS-CoV-2, Beta, Gamma or Omicron. Four days after challenge hamsters were euthanized, and infectious SARS-CoV-2 titer in lung homogenates and nasal cavity were evaluated by TCID50 measurement **(A and D)**. Lung and nasal cavity were examined for lesions by histopathological scoring and presence of viral antigen by immunohistochemistry **(B, C and E, F; Table S3-S5)**. The mean values ± SEM of all data points were shown.

Overall, the ACE2-blocking 87G7 exhibits broad and potent neutralizing activity and protects against challenge with ancestral SARS-CoV-2 and key variants of concern, including Omicron that is on its way to become the dominant lineage worldwide.

Mutations in Omicron have reshaped the antigenic landscape of the spike, likely forming a new antigenic cluster relative to all preceding VOCs and VOIs (*16, 18–22*). These mutations caused a substantial reduction of neutralizing activity of sera from Pfizer or AstraZeneca vaccine recipients and totally or partially escapes neutralization by potent neutralizing mAbs including most monoclonal antibodies that obtained emergency use authorization (*16, 18–22*). Typically, the most potent neutralizing antibodies have epitopes that overlap with the ACE2 interaction site and inhibit infection of the prototypic SARS-CoV-2 through ACE2 receptor blockage. However, these antibodies appeared significantly restricted in binding breadth due to the marked genetic diversity of the ACE2-interaction site among SARS-CoV-2 variants. The 87G7 antibody appears to be among the few exceptions of ACE2-blocking mAbs that retain potent neutralization against SARS-CoV-2 variants including Alpha, Beta, Gamma, Delta and Omicron (*5*).

Whereas effective SARS-CoV-2-specific mAb treatment for hospitalized patients has remained elusive, clinical success has been obtained in the treatment of outpatients with mild or moderate COVID-19 with anti-SARS-CoV-2 monoclonal antibodies (*36, 37*). In addition to therapeutic treatment, the development of these monoclonal antibodies may also be of value for preventive treatment of seronegative individuals including those that do not make endogenous antibodies in response to either vaccination or infection (*38*). The antigenic evolution of SARS-CoV-2 has posed a formidable challenge to the development of monoclonal antibodies for the treatment and prevention of COVID-19. While neutralization potential has been the first selection criterium of anti-SARS-CoV-2 antibody candidates for clinical use, the antibody potential for cross-neutralization through targeting highly conserved sites on the spike protein has now become much more relevant, to mitigate the risk of antibody escape by future emerging variants. Our work may contribute to the development of sustainable mAb strategies against COVID-19 using (combinations of) broadly neutralizing antibodies that are more resilient to SARS-CoV-2 antigenic diversity.

## Materials and Methods

### Viruses and cells

Calu-3 cells were maintained in Opti-MEM I (1) + GlutaMAX (Gibco) supplemented with 10% FBS, penicillin (100 IU/mL), and streptomycin (100 IU/mL) at 37°C in a humidified CO2 incubator. HEK-293T cells were cultured in DMEM supplemented with 10% FCS, sodium pyruvate (1 mM, Gibco), non-essential amino acids (1×, Lonza), penicillin (100 IU/mL), and streptomycin (100 IU/mL) at 37°C in a humidified CO2 incubator. Cell lines tested negative for mycoplasma. SARS-CoV-2 isolates were grown to passage 3 on Calu-3 cells. For stock production, infections were performed at a multiplicity of infection (moi) of 0.01 and virus was collected at 72 hours post-infection, clarified by centrifugation and stored at -80°C in aliquots. All work with infectious SARS-CoV-2 was performed in a Class II Biosafety Cabinet under BSL-3 conditions at Erasmus Medical Center. Viral genome sequences were determined using Illumina deep-sequencing as described before (*39*). The 614G virus (clade B; isolate Bavpat-1; European Virus Archive Global #026 V-03883) passage 3 sequence was identical to the passage 1 (kindly provided by Dr. Christian Drosten). The Alpha (B.1.1.7; MW947280), Gamma (P.1; OM442897), Delta (B.1.617.2; OM287123), Omicron BA.1 (B.1.1.529.1; OM287553), Omicron BA.2 (B.1.1.529.2), Lambda (C.37) and Mu (B.1.621) variant passage 3 sequences were identical to the original respiratory specimens. For Omicron, the S1 region of spike was not covered well due to primer mismatches. Therefore, the S1 region of the original respiratory specimen and passage 3 virus were confirmed to be identical by Sanger sequencing. The Beta variant (B.1.351; OM286905) passage 3 sequence contained two mutations compared the original respiratory specimen: one synonymous mutations C13860T (Wuhan-Hu-1 position) in ORF1ab and a L71P change in the E gene (T26456C, Wuhan-Hu-1 position). No other minor variants >40% were detected. SARS-CoV-2 variants of concern/interest used contained the following spike changes relative to the Wuhan-Hu-1 strain: Alpha (B.1.1.7), Δ69-70, Δ144, N501Y, A570D, D614G, P681H, T716I, S982A, D1118H; Beta (B.1.351), L18F, D80A, D215G, Δ241-243, K417N, E484K, N501Y, D614G, A701V; Gamma (P.1), L18F, T20N, P26S, D138Y, R190S, K417T, E484K, N501Y, D614G, H655Y, T1027I, V1176F; Delta (B.1.617.2), T19R, G142D, E156G, Δ157-158, L452R, T478K, D614G, P681R, D950N; Omicron BA.1 (B.1.1.529.1), A67V, Δ69-70, T95I, G142D, Δ143-145, N211I, Δ212, ins214EPE, G339D, S371L, S373P, S375F, K417N, N440K, G446S, S477N, T478K, E484A, Q493R, G496S, Q498R, N501Y, Y505H, T547K, D614G, H655Y, N679K, P681H, N764K, D796Y, N856K, Q954H, N969K, L981F; Omicron BA.2 (B.1.1.529.2), T19I, L24S, Δ25/27, G142D, V213G, G339D, S371L, S373P, S375F, T376A, D405N, R408S, K417N, N440K, S477N, T478K, E484A, Q493R, Q498R, N501Y, Y505H, D614G, H655Y, N679K, P681H, N764K, D796Y, Q954H, N969K; Lambda (C.37), G75V, T76I, R246N, Δ247-253, L452Q, F490S, D614G, T859N; Mu (B.1.621) T95I, Y144S, Y145N, R346K, E484K, N501Y, D614G, P681H, D950N.

### Expression and purification of SARS-CoV-2 S proteins

Human codon-optimized gene was synthesized at Genscript encoding the 6P-stabilized SARS-CoV-2 S ectodomain expression construct (26) (S protein residues 1–1,213, Wuhan-Hu-1 strain: GenBank: QHD43416.1) with a C-terminal T4 foldon trimerization motif followed by an octa-histidine tag and a Twin-Strep-tag® (*40*). Constructs encoding S1 (residues 1–682), the N-terminal domain (NTD, residues 1–294) or receptor-binding domain (RBD, residues 329–538) of SARS-CoV-2 S (Wuhan-Hu-1 strain), C-terminally tagged with Strep-tag have been described before (*41*). Human codon-optimized genes were synthesized encoding S1 proteins of Alpha (B.1.1.7), Beta (B.1.351), Gamma (P.1), Delta (B.1.617.2) and Omicron (B.1.1.529) VOCs described above, including a C-terminal Strep-tag. All proteins were expressed transiently in HEK-293T (ATCC® CRL-11268™) cells from pCAGGS expression plasmids, and secreted proteins were purified from culture supernatants using streptactin beads (IBA) following the manufacturer’s protocol. Spike variants with single-site residue substitutions were generated using Q5® High-fidelity DNA polymerase (NEB)-based site-directed mutagenesis.

### Immunization, hybridoma culturing and production of (recombinant) monoclonal antibodies

Harbour H2L2 mice were immunized using heterologous DNA/protein immunization protocol 16-512-22 under animal license (AVD101002016512) approved by CCD (Dutch Central Comity for animal experimentation). Mice were housed in SPF facility with cage enrichment, light switched on at 7:00 and switched off at 19:00 and with humidity at around 40%. Both female and male H2L2 mice were used. The female mice were housed up to 4 per individually ventilated cage (IVC), while males were in separate IVC cages to prevent fighting. Food was standard and water and food intake ad libitum. Mice were immunized intradermally three times bi-weekly with 50 micrograms of plasmid DNA encoding the Wuhan-Hu-1 SARS-CoV-2 S ectodomain trimer in 20 microliters of water, using the AgilePulse Intradermal electroporator system (BTX) according to the manufacturer instructions. After priming with DNA, immunization was continued in bi-weekly intervals by subcutaneous and intraperitoneal injection of 20-30 μg of antigen preparations formulated with Ribi Adjuvant System (RAS, Sigma) according to manufacturer instructions, alternating between the S ectodomain trimer and RBD of Wuhan-Hu-1 SARS-CoV-2 as antigens. Antigen specific antibody titres were monitored during immunization by taking blood samples from the mice and performing antigen-specific ELISA. High-titre mice were euthanized three to five days after the last protein boost (5 in total), B cells were collected from lymphoid tissues (lymph nodes and spleen), and hybridomas were generated by standard method using SP 2/0 myeloma cell line (ATCC #CRL-1581) as a fusion partner. Supernatants from 96 well plates (estimated to have 1-4 hybridoma clones per well) were screened for SARS-CoV-2 S binding antibodies by ELISA and neutralizing antibodies using pseudovirus neutralization assay. Selected hybridomas were subcloned by limited dilution and retested in ELISA and pseudovirus assay.

Production of recombinant human antibodies using HEK-293T was described previously (*42*). Gene blocks encoding the variable heavy (VH) and light (VL) chain sequences of 87G7 and of benchmark monoclonal antibodies REGN10933, REGN10987 (PDB ID: 6XDG) (*43*), S309 (PDB ID: 6WPS) (*44*), CR3022 (GenBank accession numbers: DQ168569.1 and DQ168570.1) (*45*), 47D11 (GenBank accession numbers: MW881223.1 and MW881224.1) (*41*) were synthesized. VH and VL sequences were separately cloned into the expression plasmids with human IgG1 heavy chain and human kappa chain constant regions, respectively using the HBM vectors pHBM 000254 (VH into pTT5-mIGK-hIgG1_HCv2) and HBM 000265 (VK into pTT5mIgK-hIgG_KCv2). Recombinant human antibodies were expressed in HEK-293T cells following transient transfection with pairs of the IgG1 heavy and light chain expression plasmids. Recombinant antibodies were purified using Protein A Sepharose (IBA) according to the manufacturer’s instructions.

### ELISA analysis of antibody binding to SARS-CoV-2 S antigens

Purified S antigens (1µg/ml) were coated onto 96-well NUNC Maxisorp plates (Thermo Scientific) at room temperature (RT) for 3 h followed by three washing steps with Phosphate Saline Buffer (PBS) containing 0.05% Tween-20. Plates were blocked with 3% bovine serum albumin (BSA, Fitzgerald) in PBS with 0.1% Tween-20 at 4℃ overnight. 87G7 mAb was allowed to bind to the plates at 5-fold serial dilutions, starting at 10 μg/ml diluted in PBS containing 3% BSA and 0.1% Tween20, at RT for 1 h. Antibody binding to the S proteins was determined using a 1:2000 diluted HRP conjugated goat anti-human IgG, (ITK Southern Biotech) for 1 h at RT and tetramethylbenzidine substrate (BioFX). Readout for binding was done at 450 nm (OD450) using the ELISA plate reader (EL-808, Biotek).

### Antibody binding kinetics and affinity measurement

87G7 (21 nM) was loaded onto Protein A biosensors (ForteBio) for 10 min. Antigen binding was performed by incubating the biosensor with 2-fold dilutions of recombinant SARS-CoV-2 S1 monomer or S ectodomain trimer for 10 min followed by a long dissociation step (30 min) to observe the decrease of the binding response. The affinity constant K_D_ was calculated using 1:1 Langmuir binding model on Fortebio Data Analysis 7.0 software.

### Biolayer interferometry-based binding competition assay

Binding competition was performed using biolayer interferometry (Octet Red348; ForteBio), as described previously (*41, 42*). In brief, SARS-CoV-2 S ectodomain trimer (50 μg/ml) was immobilized onto the anti-strep mAb-coated protein A biosensor. After a brief washing step, the biosensor tips were immersed into a well containing primary mAb (50 μg/ml) for 15 min and subsequently into a well for 15 min containing the competing mAb (secondary mAb; 50 μg/ml) or recombinant soluble ACE2. A 3 to 5-min washing step in PBS was included in between steps.

### ELISA-based receptor-binding inhibition assay

The ACE2 receptor-binding inhibition assay was performed as described previously (*41, 42*). Recombinant soluble ACE2 was coated on NUNC Maxisorp plates (Thermo Scientific) at 1µg/well at RT for 3 h. Plates were washed three times with PBS containing 0.05% Tween-20 and blocked with 3% BSA (Fitzgerald) in PBS containing 0.1% Tween-20 at 4 °C overnight. Recombinant SARS-CoV-2 S RBD domain (200 nM) and serially diluted mAbs were mixed and incubated for 2 h at RT. The mixture was added to the plate for 2 h at 4 °C, after which plates were washed three times. Binding of SARS-CoV-2 S RBD domain to ACE2 was detected using 1:2000 diluted HRP-conjugated anti-StrepMAb (IBA) that recognizes the Strep-tag affinity tag on the SARS-CoV-2 S RBD domain. Detection of HRP activity was performed as described above (ELISA section).

### Pseudovirus neutralization assay

Human codon-optimized genes encoding the spike proteins of SARS-CoV-2 S proteins corresponding to ancestral Wuhan-Hu-1 virus (Genbank: NC_045512.2) or variants of concern Alpha (B.1.1.7), Beta (B.1.351), Gamma (P.1), Delta (B.1.617.2) and Omicron (B.1.1.529) were synthesized by GenScript. The production of SARS-CoV-2 S pseudotyped vesicular stomatitis virus (VSV) and the neutralization assay were performed as described previously (*41*). In brief, HEK-293T cells at 70∼80% confluency were transfected with the pCAGGS expression vectors encoding SARS-CoV-2 S with a C-terminal cytoplasmic tail 18-residue truncation to increase cell surface expression levels. Cells were infected with VSV G pseudotyped VSVΔG bearing the firefly (*Photinus pyralis*) luciferase reporter gene at 48 hours after transfection. Twenty-four hours later, the supernatant was harvested and filtered through 0.45 μm membrane. Pseudotyped VSV was titrated on VeroE6 cells. In the virus neutralization assay, 3-fold serially diluted mAbs were pre-incubated with an equal volume of virus at RT for 1 h, and then inoculated on VeroE6 cells, and further incubated at 37°C. After 20 h, cells were washed once with PBS and lysed with Passive lysis buffer (Promega). The expression of firefly luciferase was measured on a Berthold Centro LB 960 plate luminometer using D-luciferin as a substrate (Promega). The percentage of neutralization was calculated as the ratio of the reduction in luciferase readout in the presence of mAbs normalized to luciferase readout in the absence of mAb. The half maximal inhibitory concentrations (IC50) were determined using 4-parameter logistic regression (GraphPad Prism v8.3.0).

### Live virus neutralization assay

Human monoclonal antibodies were tested for live virus neutralization using a plaque reduction neutralization (PRNT) assay. PRNT was performed according to a previously published protocol (*39*), with minor modifications. Briefly, 50 μl of serially diluted antibody in Opti-MEM I (IX) + GlutaMAX (Gibco, USA) was mixed 1:1 with virus (400 PFU) and incubated at 37°C for 1 hour before layering over fully confluent monolayers of Calu-3 cells (washed once prior with Opti-MEM I (IX) + GlutaMAX). After 8 h of infection, the cells were fixed with formalin, permeabilized with 70% ethanol, washed in PBS and stained using rabbit anti-SARS-CoV nucleocapsid (SinoBiological, 1:2000 in 0.1% bovine serum albumin (BSA) in PBS) followed by goat anti-rabbit Alexa Fluor 488 antibody (Invitrogen, 1:2000 in 0.1% BSA in PBS). Plates were scanned on the Amersham Typhoon Biomolecular Imager (GE Healthcare, USA). Data were analyzed using ImageQuantTL 8.2 image analysis software (GE Healthcare). The PRNT titer was calculated using Graphpad Prism 9, calculating a 50% reduction in infected cells counts based on non-linear regression with bottom constraints of 0% and top constraints of 100%.

### Cryo-electron microscopy sample preparation and data collection

The 87G7 Fab fragment was digested from the IgG with papain using a Pierce Fab Preparation Kit (Thermo Fisher Scientific), according to the manufacturer’s instructions. Spike-Fab complexes were prepared under two conditions. For the first condition, 4 μl of SARS-CoV-2 hexaproline spike ectodomain, at a concentration of 28 μM (based on the molecular weight of the spike protomer) was combined with 1 μl of 150 μM 87G7 Fab and incubated for ∼10 min at RT before blotting and plunge freezing. For the second condition, 3.5 μl of 28 μM SARS-CoV-2 hexaproline spike ectodomain was combined with 1 μl of 150 μM 87G7 Fab and then incubated for ∼10 min at RT. Immediately before blotting and plunge freezing, 0.5 μl of 0.2% *(w/v)* fluorinated octyl maltoside (FOM) was added to the sample, resulting in a final FOM concentration of 0.02% *(w/v)*. For both conditions, 3 μl of spike-Fab complex was applied to glow-discharged (20 mAmp, 30 sec, Quorum GloQube) Quantifoil R1.2/1.3 grids (Quantifoil Micro Tools GmbH), blotted for 5 s using blot force 2 and plunge frozen into liquid ethane using Vitrobot Mark IV (Thermo Fisher Scientific). The data were collected on a Thermo Scientific™ Krios™ G4 Cryo Transmission Electron Microscope (Cryo-TEM) equipped with Selectris X Imaging Filter (Thermo Fisher Scientific) and Falcon 4 Direct Electron Detector (Thermo Fisher Scientific) operated in Electron-Event representation (EER) mode. Data processing was performed in Relion 3.1 (*46*) and cryoSPARC™ (*47*)single particle analysis suites. Raw data were imported in cryoSPARC™. After Patch motion correction and Patch CTF estimation, 313,636 particles were picked from 1331 images from 0.02% FOM dataset and 621,175 particles were picked from 2500 images without FOM. After 2D classification and heterogenous refinement, the best particle stack consisting of 133,550 particles was subjected to non-uniform refinement (*48*) with C3 symmetry imposed yielding a Spike-Fab complex cryo-EM map with an overall resolution of 2.9 Å. Following global refinement, a soft mask encompassing one RBD with the Fab bound was made in UCSF Chimera (*49*). Particles were imported into Relion 3.1 and, using the “relion_particle_symmetry_expand” tool, each particle from the C3-symmetry–imposed reconstruction was assigned three orientations corresponding to its symmetry related views. The soft mask was placed over a single RBD-Fab region of the map, and the symmetry-expanded particles were subjected to masked 3D classification without alignment using a regularization parameter (‘T’ number) of 20. Following a single round of focused classification, the best particle stack consisting of 72,118 particles was imported back to cryoSPARC™ and refined without imposing symmetry using the local refinement job, yielding a map with a global resolution of 4.9 Å. The nominal resolutions and local resolution estimations for the global and local refinements were performed in Relion 3.1. The ‘Gold Standard’ Fourier shell correlation (FSC) criterion (FSC = 0.143) was used for resolution estimates. Finally, the globally and locally refined maps were masked and sharpened using DeepEMhancer tool (*50*), as implemented in COSMIC2 (*51*), and combined using the “vop add” command in UCSF Chimera (*49*). Data collection and reconstruction parameters can be found in Table 1.

### Model building and refinement

UCSF Chimera (*49*) (version 1.15.0) and Coot (*52*) (version 0.9.6) were used for model building. As a starting point for modelling the 87G7-bound spike, the crystal structure of the SARS-CoV-2-S N-terminal domain (residues 14-308; PDB ID: 7B62 (*53*)), the fully open SARS-CoV-2-S model (residues 309-332 and 527-1145; PDB ID: 7K4N (*29*)) and RBD crystal structure (residues 333-526; PDB ID 6M0J (*54*)) were individually rigid-body fitted into the composite density map using the UCSF Chimera “Fit in map” tool (*49*). Subsequently, the models were combined, and the peptide sequence was adjusted to match the 6P spike construct used in this study. For modelling the 87G7 Fab fragment, atomic coordinates of the heavy chain (HC) and the light chain (LC) variable regions were generated using the phyre2 server (*55*) and rigid body fitted into the EM density map using the UCSF Chimera ‘fit in map’ tool and then combined with the spike model. The resulting model was then edited in Coot using the ‘real-space refinement (*52*), carbohydrate module (*56*) and ‘sphere refinement’ tool. Iterative rounds of manual fitting in Coot and real space refinement in Phenix (*57*) were carried out to improve non-ideal rotamers, bond angles and Ramachandran outliers. During refinement with Phenix, secondary structure and non-crystallographic symmetry restraints were imposed. The final model was validated with MolProbity (*58*), EMRinger (*59*) and Privateer (glycans) (*60, 61*).

### Analysis and visualization

Spike residues interacting with 87G7 were identified using PDBePISA (*62*) and LigPlot^+^ (*63*) . Surface coloring of the SARS-CoV-2 RBD according to sequence conservation and the Kyte-Doolittle hydrophobicity scale was performed in UCSF ChimeraX (*64*). The UCSF Chimera “MatchMaker” tool was used to obtain RMSD values, using default settings. Figures were generated using UCSF Chimera (*49*) and UCSF ChimeraX (*64*). *Structural biology applications used in this project were compiled and configured by SBGrid* (*65*).

### Mouse challenge experiment

*In vivo* prophylactic and therapeutic efficacy of mAb 87G7 against challenge with SARS-CoV-2 and four variants of concern, was evaluated in heterozygous K18-hACE2 C57BL/6J mice (strain: 2B6.Cg-Tg(K18-ACE2)2Prlmn/J) obtained from The Jackson Laboratory. Groups of 14-week-old mice (n = 5), were given 200 μg of 87G7 or isotype control antibody (equivalent to 10 mg of the antibody per kg) by intraperitoneal injection, 16 h before or one day after intranasal inoculation with a lethal dose of the indicated SARS-CoV-2 strain (10^5^ PFU/mouse). Virus inoculations were performed under anesthesia that was induced with isoflurane, and all efforts were made to minimize animal suffering. All animals were housed in a self-contained ventilated rack (Tecniplast, IT), with the light switched on at 7:30 and switched off at 19:30. The ambient temperature was 19.5-22 °C and with humidity at 35-40%. Animal protection studies were carried out under the animal permit PROEX-146.6/20, approved by the Community of Madrid (Spain), and performed in biosafety level 3 facilities at CISA-INIA (Madrid).

To quantify infectious SARS-CoV-2 virus particles, one fourth of the right lung was homogenized using a MACS homogenizer (Miltenyi Biotec) according to the manufacturer’s protocols. Virus titrations were done using plaque assay performed on Vero E6 cells following standard procedures. In brief, cells were overlaid with DMEM containing 0.6% low-melting agarose and 2% FBS, fixed with 10% formaldehyde and stained with 0.1% crystal violet at 72 h post-infection.

To quantify viral antigen by immunohistochemistry, left lung lobes were fixed in 10% buffered formalin (Chemie Vertrieb GmbH & Co Hannover KG, Hannover, Germany). Left lung lobes were pre-fixed by injections of 10% buffered formalin as recommended by Meyerholz et al (*66*) to ensure an optimal histopathological evaluation **(Table S2)**.

### Hamster challenge experiment

During the experiment, the animals were under veterinary observation and all efforts were made to minimize distress. Approval for the experiments was given by the German Niedersächsisches Landesamt für Verbraucherschutz und Lebensmittelsicherheit (LAVES file number 21/3755) and by the Dutch authorities (Project license number 27700202114492-WP12). Syrian hamsters (*Mesocricetus auratus*, 6-10 weeks old) were housed under BSL-3 conditions, starting 10 days prior to the experiment. 87G7 or a non-SARS-CoV-2 human IgG control antibody were injected intraperitoneally in a volume of 500 µl. The hamsters were challenged intranasally, 24 h after or 12 h before antibody inoculation, with 10^4^ TCID50 of the respective SARS-CoV-2 variants, respectively. The animals were monitored for body weight loss and clinical symptoms twice daily until they were humanely euthanized four days after infection. Antibody injection, with challenge virus and euthanasia were performed under isoflurane anesthesia. Left nasal turbinates and left lung lobe were fixed in 10% buffered formalin (Chemie Vertrieb GmbH & Co Hannover KG, Hannover, Germany) from the investigated hamsters. Left lung lobes were pre-fixed by injections of 10% buffered formalin as recommended by Meyerholz et al. (*66*) to ensure an optimal histopathological evaluation. Left nasal turbinates, following formalin fixation, were decalcified in soft tissue decalcifier (Roth # 6484.2) for about 14 days prior to routine tissue processing.

To quantify infectious SARS-CoV-2 virus particles, lung and nasal turbinate tissues were homogenized using a TissueLyser II (Qiagen) and infectious SARS-CoV-2 virus particles in tissue homogenates were quantified on Vero E6 cells. Cells were infected with 10 fold serial dilutions of the homogenized tissue prepared in DMEM + 2 % FBS (starting dilution 100- and 10-fold for lung and nasal turbinate homogenate, respectively). Plates were further incubated in a humidified atmosphere, at 37°C, 5% CO2. Cytopathic effect was evaluated 5 days post infection. Omicron samples were titrated in Calu-3 cells due to the low infectivity of Omicron in Vero cells. In this case, after 5 day incubation, cells were fixed with 4% PFA and stained using an anti-SARS-CoV-2 nucleocapsid antibody (Sinobiological). Virus titers (TCID50/ml) were calculated using the Spearman-Karber method.

Formalin-fixed, paraffin-embedded (FFPE) tissue was used for histology and immunohistochemistry. Histopathological lesions were evaluated on hematoxylin-eosin (HE) stained sections. For the detection of viral antigen in Syrian golden hamsters, immunohistochemistry with a monoclonal antibody detecting SARS-CoV/SARS-CoV-2 nucleocapsid (Sino Biological 40143-MM05) was performed on FFPE tissue sections, as described previously (*67, 68*). Briefly, tissue sections were dewaxed and rehydrated, followed by endogenous peroxidase blocking for 30 min at RT. Antigen retrieval was performed in Na_2_H_2_EDTA buffer for 20 minutes in a microwave at 800 W. The primary antibody (dilution 1:4000) was applied for 1 h at RT. Sections were subsequently rinsed, and secondary labeling was performed using the respective peroxidase-labeled polymer (Dako Agilent Pathology Solutions, K4003) for 30 min for 60 min at RT. Visualization of the reaction was accomplished by incubation in chromogen 3,3-diaminobenzidine tetrahydrochloride (DAB, 0.05%) and 0.03% H_2_O_2_ in PBS for 5 min. The slides were afterwards counterstained with Mayer’s hematoxylin for 1 min. Nasal turbinates were evaluated on a full-length longitudinal section of the nose including respiratory and olfactory epithelium. Assessment of histopathological lesions in the nasal turbinates was performed with a semi-quantitative score system, as described previously with minor modifications . Quantification of the viral antigen in the nasal epithelium was performed using a semi-quantitative score. Hamsters left lung lobe was evaluated on one cross-section (at the level of the entry of the main bronchus) and one longitudinal section (along the main bronchus) of the entire left lung lobe. Assessment of histopathological lesions and viral load in the lung was performed with a semi-quantitative scoring system, as described previously with minor modifications (*69*). System for semi-quantitative scoring of histopathological lesions and viral antigen in nose and lung is shown in **Table S3-S5**. Histopathological semi-quantitative evaluations were performed by veterinary pathologists (GB, MC, FA) and subsequently confirmed by a European board certified veterinary pathologist (WB). During the evaluation, the pathologist was blinded regarding the treatment groups and used virus strains.

## Acknowledgments

We thank Caroline Schütz, Julia Baskas, Jana-Svea Harre, Vera Nijman, Marianthi Chatziandreou and Rutger Brouwer for technical support. This study was done within the framework of the Utrecht Molecular Immunology Hub - Utrecht University and the research programme of the Netherlands Centre for One Health (www.ncoh.nl). Funding: The MANCO project has received funding from the European Union’s Horizon 2020 research and innovation programme under grant agreement No 101003651). This work made use of the Dutch national e-infrastructure with the support of the SURF Cooperative using grant no. EINF-2453. This research was funded by the Deutsche Forschungsgemeinschaft (DFG; German Research Foundation) - 398066876/GRK 2485/1; BMBF (Federal Ministry of Education and Research) project entitled RAPID (Risk assessment in re-pandemic respiratory infectious diseases), 01KI1723G, Ministry of Science and Culture of Lower Saxony in Germany (14 - 76103-184 CORONA-15/20)

## Author contributions

Gene cloning, protein expression and purification, WD, JL, TS, RvH and DD; immunization, hybridoma fusion and screening, subcloning, sequencing, production and purification: RvH and DD; affinity measurements, epitope binning and neutralization assays: WD, JL, AM and MML; cryo-EM grid preparation and data collection, ID; cryo-EM data processing, atomic modelling and interpretation, DLH; animal experiments, MGH, FK and GvA; pathological investigation, FA., GB, MC, WB; supervision, DLH, FJMvK, BLH, LE, ADMEO, FG, and BJB; study conception and coordination, FG and BJB; manuscript writing, WD, DLH and BJB, with input from all other authors.

## Data and materials availability

The globally and locally refined cryo-EM maps have been deposited to the Electron Microscopy Data Bank under the accession codes EMD-14250 and EMD-14271, respectively. The atomic model of the 87G7-bound spike has been deposited to the Protein Data Bank under the accession code 7R40. Materials generated in this study are available on reasonable request.

## Supplementary Figures and Tables

**Figure S1.**
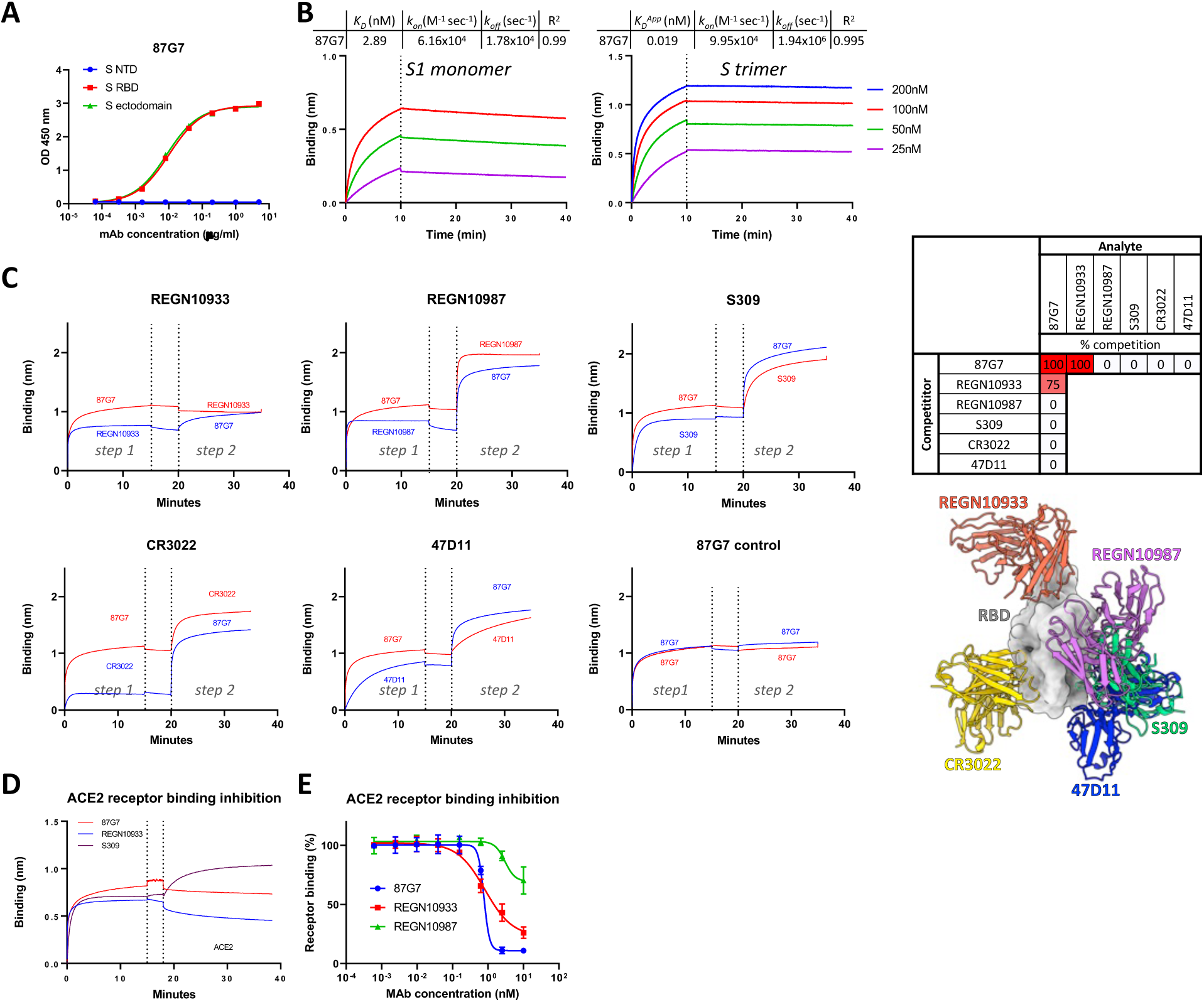
87G7 epitope binning and ACE2 binding inhibition. **(A)** ELISA binding curves of 87G7 to the plate-immobilized N-terminal domain (NTD), the receptor-binding domain (RBD) or ectodomain of SARS-CoV-2 S. **(B)** Binding kinetics of 87G7 to SARS-CoV-2 S measured by biolayer interferometry (BLI). 87G7 mAb was loaded at optimal concentration (21 nM) onto the anti-human Fc biosensor for 10 mins, after which association of antigen was achieved by incubating the sensor with a 2-fold dilution series of recombinant SARS-CoV-2 S1 monomer or S ectodomain trimer for 10 min, followed by a dissociation step in PBS for 30 min. *K_D_* : equilibrium dissociation constant. *k_on_* : association rate constant, *k_off_* : dissociation rate constant. *K_D_^App^* reflects the ‘apparent affinity’ between IgG antibodies and spike trimer. **(C)** Binding competition between 87G7 and benchmarking antibodies to SARS-CoV-2 S ectodomain trimer evaluated using BLI. The strep-tagged SARS-CoV-2 S antigen was loaded to the anti-human Fc biosensor bound with antibodies against Strep tag. The competitor antibody was bound to spike (step 1) before incubation with the analyte antibody (step 2) as indicated, and percent competition bins are indicated in the table (dark red: >90% competition, light red; 40-80% competition, white: <10% competition). Data from a representative experiment (out of two) are shown. Benchmarking antibodies tested for binding competition with 87G7 are shown in complex with SARS-CoV-2 RBD and include REGN10933 (PDB: 6XDG), REGN10987 (PDB: 6XDG), S309 (parent of VIR-7831, PDB: 6WPS), CR3022 (PDB: 6W41) and 47D11 (PDB: 7AKD) **(D)** BLI-based receptor-binding inhibition assay. SARS-CoV-2 S ectodomain bound to the sensor was pre-incubated with 87G7 or two control mAbs REGN10933 (ACE2 binding competitor) or S309 (non-ACE2 competing), followed by a washing step and subsequent exposure to soluble human ACE2 receptor. The experiment was performed twice, data from a representative experiment is shown. **(E)** ELISA-based receptor-binding inhibition assay. SARS-CoV-2 S ectodomain pre-incubated with serially diluted 87G7 or two control mAbs REGN10933 or REGN10987 (both ACE2 binding competitors) was added to ELISA plates coated with soluble human ACE2. Spike binding to ACE2 was detected using an HRP-conjugated antibody recognizing the C-terminal Strep-tag on SARS-CoV-2 S ectodomain. Data points represent the average ± SDM, for n = 3 replicates from two independent experiments.

**Figure S2.**
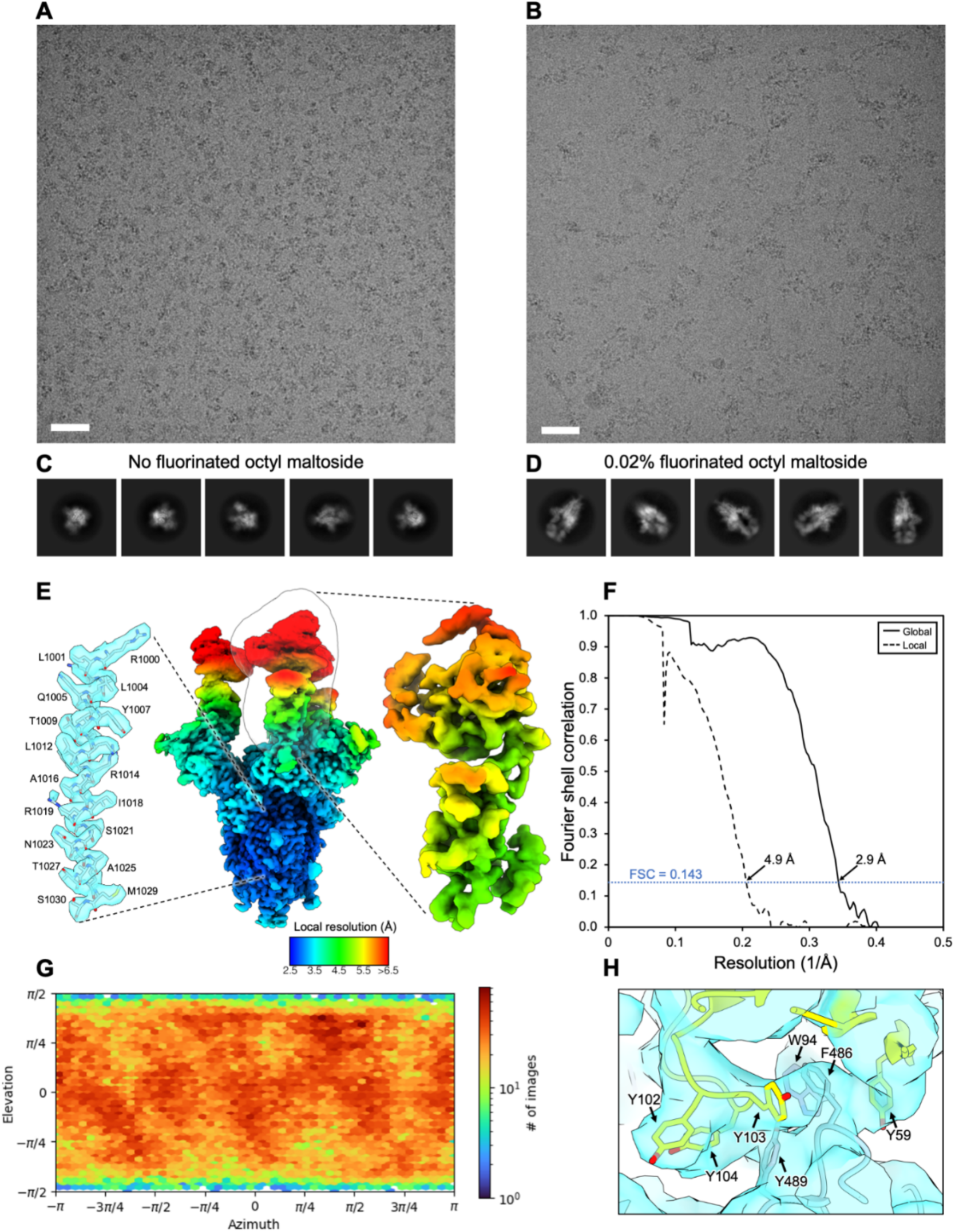
Cryo-EM data processing of SARS-CoV-2 S bound to 87G7 Fab. **(A)** Representative motion-corrected micrograph of the 87G7-bound SARS-CoV-2 spike ectodomains embedded in vitreous ice. Scale bar = 50 nm. **(B)** As shown in A, for the sample incubated with 0.2% fluorinated octyl maltoside. **(C)** Representative reference-free 2D class averages generated in cryoSPARC. **(D)** As shown in C, for the sample incubated with 0.2% fluorinated octyl maltoside. **(E)** DeepEMhancer filtered EM density maps for the globally refined spike-87G7 complex and locally refined RBD-87G7 complex, colored according to local resolution which was calculated in Relion3.1. The outline of the local refinement mask is overlaid with the globally refined map. Representative density and fitted atomic coordinates for the S2 region of the globally refined map is shown on the left. **(F)** Gold-standard Fourier shell correlation (FSC) curves generated from the independent half maps contributing to the 2.9 Å resolution global refinement and 4.9 Å resolution local refinement. **(G)** Angular distribution calculated in cryoSPARC for particle projections in the local refinement. **(H)** Cryo-EM density for the locally refined 87G7 epitope-paratope region with the fitted atomic coordinates. RBD residues are colored blue and the light- and heavy-chain variable domains are colored purple and yellow, respectively.

**Figure S3.**
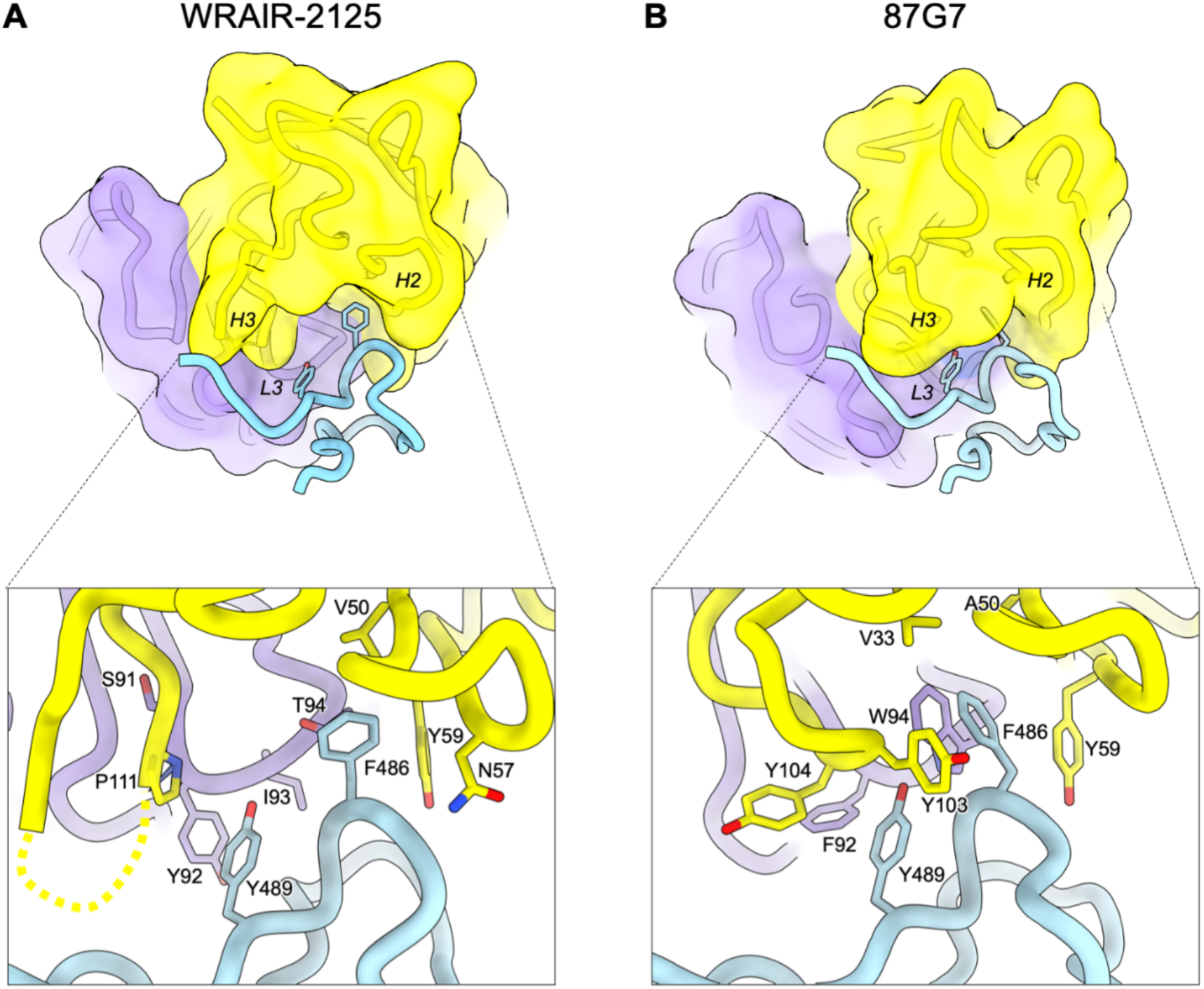
Structural comparison of WRAIR-2125 and 87G7. **(A)** The top panel shows a surface representation of the WRAIR-2125 Fab fragment variable domains, rendered at 6 Å resolution, in complex with SARS-CoV-2 RBD residues 470-494 (generated using PDB ID: 7N4L). The CDR H2, H3 and L3 loop are labelled in italics. The Fab heavy and light chain are colored yellow and purple and RBD residues are colored blue. The bottom panel shows a close-up view showing selected interactions formed between WRAIR-2125 residues and the SARS-CoV-2 RBD residues Y489 and F489. The unmodelled region of the CDR H3 loop is indicated with a dashed line. **(B)** As shown in A for 87G7.

**Table S1.**
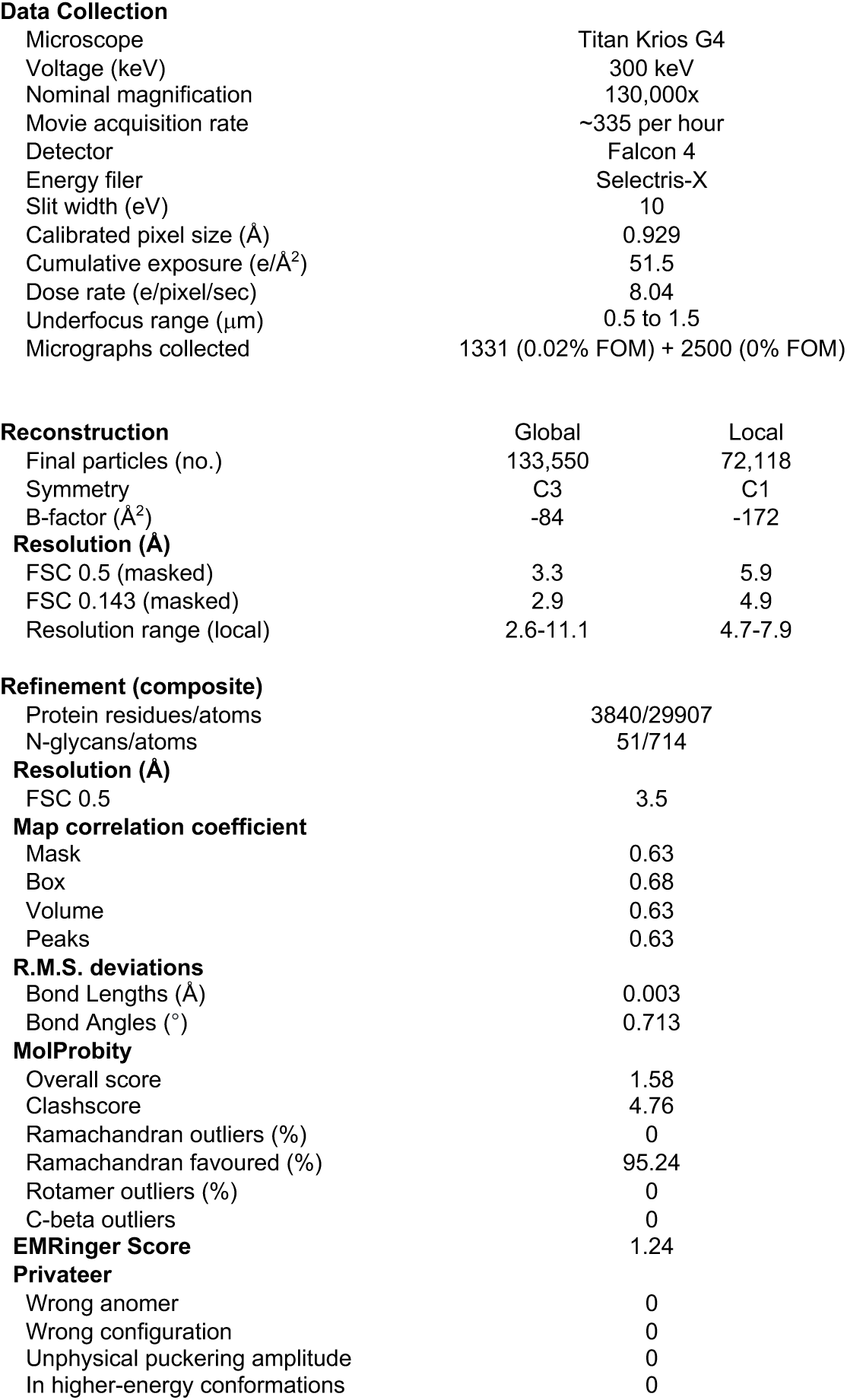
Summary of data acquisition and image processing statistics. Related to Figure 2 and 3.

**Table S2.** Scoring system of mice lung SARS-CoV antigen immunohistochemistry. Related to Figure 4B and 4E.

|  |  |  |
| --- | --- | --- |
| **1. Immunohistochemistry** |  |  |
| **1.1 Extent alveolar antigen** |  |  |
| 0 | No | No specific signal |
| 1 | Minimal | Single small foci with immunolabelled pneumocytes, less than 1 % of alveoli |
| 2 | Mild | Multifocal areas with immunolabelled pneumocytes, 2-25 % of alveoli |
| 3 | Moderate | Multifocal areas with immunolabelled pneumocytes, 26-50 % of alveoli |
| 4 | Marked | Multifocal areas with immunolabelled pneumocytes, 51-75 % of alveoli |
| 5 | Subtotal | Multifocal areas with immunolabelled pneumocytes, > 75 % of alveoli |
| **1.2 Extent airway (bronchi and bronchioli) antigen** |  |  |
| 0 | No | No specific signal |
| 1 | Minimal | Single small foci with immunolabelled epithelia, less than 1 % of airways |
| 2 | Mild | Multifocal areas with immunolabelled epithelia, 2-25 % of airways |
| 3 | Moderate | Multifocal areas with immunolabelled epithelia, 26-50 % of airways |
| 4 | Marked | Multifocal areas with immunolabelled epithelia, 51-75 % of airways |
| 5 | Subtotal | Multifocal areas with immunolabelled epithelia, > 75 % of airways |
| Total immunohistochemistry score |  |  |
| **Semiquantitative score (antigen):** Sum of alveolar and airway score |  |  |

**Table S3.** Scoring system of hamster lung (alveolar, airway and vascular) lesions, Related to Figure 5B and 5E.

|  |  |  |
| --- | --- | --- |
| **1. Alveolar lesions** |  |  |
| **1.1 Extent alveolar inflammation** |  |  |
| 0 | No | No inflammation |
| 1 | Minimal | Single small foci of mild inflammatory infiltrates, affecting max. 1% of tissue, (significance questionable) |
| 2 | Mild | 2-25 % of tissue affected |
| 3 | Moderate | 26-50 % of tissue affected |
| 4 | Marked | 51-75 % of tissue affected |
| 5 | Subtotal | > 75 % of tissue affected |
| **1.2 Severity alveolar inflammation (scored in area of maximal severity)** |  |  |
| 0 | No | No inflammatory infiltrates |
| 1 | Minimal | Few inflammatory cells in alveolar septae or lumina, alveolar architecture maintained (significance questionable) |
| 2 | Mild | Mild septal and luminal infiltrates (2-3 cells thick), alveolar architecture maintained |
| 3 | Moderate | Moderate septal and luminal infiltrates, occasionally obscuring alveolar architecture |
| 4 | Marked | Marked septal and luminal infiltrates with large areas with completely obscured alveolar architecture |
| **1.3 Alveolar damage** |  |  |
| 0/1 | No/Yes | Necrosis/desquamation/loss (denuded septae) of alveolar cells |
| 0/1 | No/Yes | Intraalveolar fibrin/hyaline membranes |
| 0/1 | No/Yes | Atypical/large or multinucleated cells lining alveoli |
| 0/1 | No/Yes | Alveolar edema |
| 0/1 | No/Yes | Alveolar hemorrhage |
| **Total alveolar lesions score** |  |  |
| **Sum of all scores** |  |  |
| **2. Airway lesions** |  |  |
| **2.1 Extent airway inflammation** |  |  |
| 0 | No | No inflammation |
| 1 | Minimal | Single small foci of mild inflammatory infiltrates, affecting max. 1% of tissue, (significance questionable) |
| 2 | Mild | 2-25 % of tissue affected |
| 3 | Moderate | 26-50 % of tissue affected |
| 4 | Marked | 51-75 % of tissue affected |
| 5 | Subtotal | > 75 % of tissue affected |
| **2.2 Severity airway (bronchi/bronchiole) inflammation (scored in area of maximal severity)** |  |  |
| 0 | No | No inflammation |
| 1 | Minimal | Rare peribronchial/peribronchiolar, mild, mononuclear infiltrates (significance questionable) |
| 2 | Mild | Mild, mononuclear and granulocytic bronchitis/bronchiolitis (with exocytosis of inflammatory cells into epithelium, occasional single cell necrosis and mild, peribronchial/peribronchiolar infiltrates) |
| 3 | Moderate | Moderate, mononuclear and granulocytic to necrotizing bronchitis/bronchiolitis (with exocytosis of inflammatory cells into epithelium, frequent single cell necrosis and intraluminal debris + moderate, peribronchial/peribronchiolar infiltrates) |
| 4 | Marked | Marked, mononuclear and granulocytic and necrotizing bronchitis/bronchiolitis with widespread exocytosis of inflammatory cells into epithelium, frequent epithelial necrosis and intraluminal debris, severe peribronchial/peribronchiolar infiltrates) |
| **Total airway lesions score** |  |  |
| **Sum of all scores** |  |  |
| **3. Vascular lesions** |  |  |
| **3.1 Extent vacuolopathy/vasculitis (large and medium sized vessels with visible tunica media, characterized by endothelialitis and mural infiltrates with disruption of vessel walls)** |  |  |
| 0 | No | No lesions |
| 1 | Minimal | Single vessels with mild inflammation (significance questionable) |
| 2 | Mild | 2-25 % of vessels affected |
| 3 | Moderate | 26-50 % of vessels affected |
| 4 | Marked | 51-75 % of vessels affected |
| 5 | Subtotal | > 75 % of vessels affected |
| **3.2 Severity perivascular infiltrates (scored in area of maximal severity)** |  |  |
| 0 | No | No inflammation |
| 1 | Minimal | Single vessels with endothelial hypertrophy and few perivascular cells (no continuous cuff, significance questionable) |
| 2 | Mild | 1-2 cell layers of perivascular cuffs |
| 3 | Moderate | 3-5 cell layers of perivascular cuffs |
| 4 | Marked | > 5 cell layers of perivascular cuffs |
| **3.3 Vascular lesions** |  |  |
| 0/1 | No/Yes | Perivascular edema |
| 0/1 | No/Yes | Perivascular hemorrhage |
| **Total vascular lesions score** |  |  |
| **Sum of all scores** |  |  |

**Semiquantitative score (lesion lung):** sum of total alveolar, airway and vascular lesions score

**Table S4:**
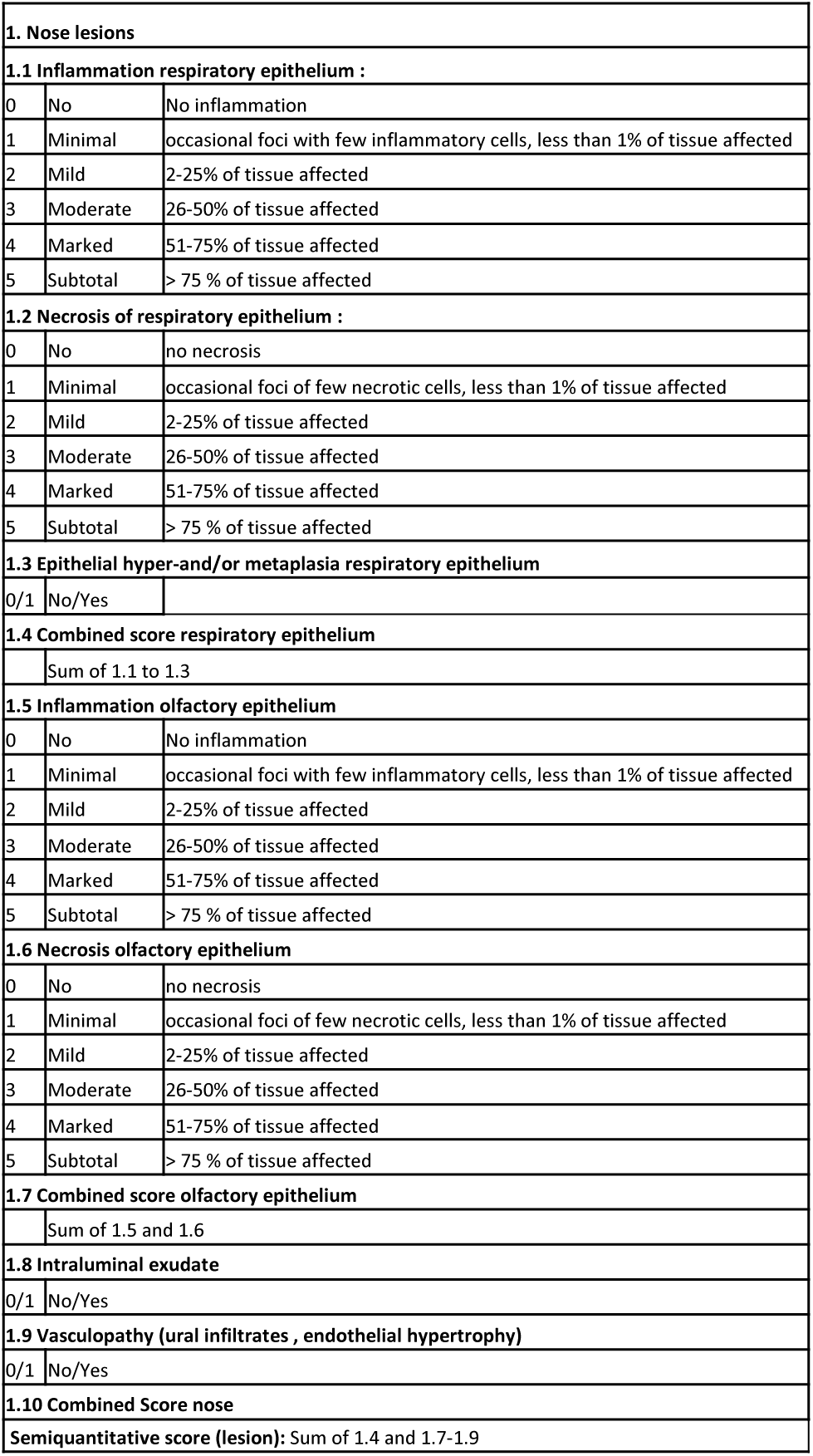
Scoring system of nasal cavity lesions. Related to Figure 5B and 5E.

**Table S5:** Scoring system of hamster lung and nose SARS-CoV-2 antigen immunohistochemistry. Related to Figure 5C and 5F.

|  |  |  |
| --- | --- | --- |
| **1. Immunohistochemistry (lung)** |  |  |
| **1.1 Extent alveolar antigen** |  |  |
| 0 | No | No specific signal |
| 1 | Minimal | Single small foci with immunolabelled pneumocytes, less than 1 % of alveoli |
| 2 | Mild | Multifocal areas with immunolabelled pneumocytes, 2-25 % of alveoli |
| 3 | Moderate | Multifocal areas with immunolabelled pneumocytes, 26-50 % of alveoli |
| 4 | Marked | Multifocal areas with immunolabelled pneumocytes, 51-75 % of alveoli |
| 5 | Subtotal | Multifocal areas with immunolabelled pneumocytes, > 75 % of alveoli |
| **1.2 Extent airway (bronchi and bronchioli) antigen** |  |  |
| 0 | No | No specific signal |
| 1 | Minimal | Single small foci with immunolabelled epithelia, less than 1 % of airways |
| 2 | Mild | Multifocal areas with immunolabelled epithelia, 2-25 % of airways |
| 3 | Moderate | Multifocal areas with immunolabelled epithelia, 26-50 % of airways |
| 4 | Marked | Multifocal areas with immunolabelled epithelia, 51-75 % of airways |
| 5 | Subtotal | Multifocal areas with immunolabelled epithelia, > 75 % of airways |
| **Total immunohistochemistry score** |  |  |
| **Semiquantitative score (antigen):** Sum of alveolar and airway score |  |  |
| **2. Immunohistochemistry nose** |  |  |
| **2.1 Respiratory Epithelium:** |  |  |
| 0 | No | No signal |
| 1 | Minimal | single positive cells, up to 1% of epithelium |
| 2 | Mild | 2-25% of epithelium positive |
| 3 | Moderate | 26-50% of epithelium positive |
| 4 | Marked | 51-75% of epithelium positive |
| 5 | Subtotal | > 75 % of epithelium positive |
| **2.2 Olfactory Epithelium:** |  |  |
| 0 | No | No signal |
| 1 | Minimal | single positive cells, up to 1% of epithelium |
| 2 | Mild | 2-25% of epithelium positive |
| 3 | Moderate | 26-50% of epithelium positive |
| 4 | Marked | 51-75% of epithelium positive |
| 5 | Subtotal | > 75 % of epithelium positive |
| **2.3 Intraluminal exudate** |  |  |
| 0/1 | No/Yes |  |
| **2.4 Subepithelial cells** |  |  |
| 0/1 | No/Yes |  |
| **2.5 Combined Score** |  |  |
| **Semiquantitative score (antigen):** Sum of all scores |  |  |

## References

1. K. Tao et al., The biological and clinical significance of emerging SARS-CoV-2 variants. Nat Rev Genet. 22, 757–773 (2021).

2. P. Wang et al., Increased resistance of SARS-CoV-2 variant P.1 to antibody neutralization. Cell Host & Microbe. 29, 747–751.e4 (2021).

3. P. Wang et al., Antibody resistance of SARS-CoV-2 variants B.1.351 and B.1.1.7. Nature (London). 593, 130–135 (2021).

4. A. Wilhelm, et al., Reduced Neutralization of SARS-CoV-2 Omicron Variant by Vaccine Sera and Monoclonal Antibodies. medRxiv., 2021.12.07.21267432 (2021).

5. E. Cameroni et al., Broadly neutralizing antibodies overcome SARS-CoV-2 Omicron antigenic shift. Nature. 602, 664–670 (2021).

6. S. Cele et al., SARS-CoV-2 Omicron has extensive but incomplete escape of Pfizer BNT162b2 elicited neutralization and requires ACE2 for infection. Nature., 1–3 (2021).

7. W. Dejnirattisai et al., Reduced neutralisation of SARS-CoV-2 omicron B.1.1.529 variant by post-immunisation serum. Lancet.(2021).

8. T. G. Caniels et al., Emerging SARS-CoV-2 variants of concern evade humoral immune responses from infection and vaccination. Sci Adv. 7, eabj5365 (2021).

9. W. F. Garcia-Beltran et al., mRNA-based COVID-19 vaccine boosters induce neutralizing immunity against SARS-CoV-2 Omicron variant. Cell. 185, 457–466.e4 (2022).

10. M. J. van Gils, et al., Four SARS-CoV-2 vaccines induce quantitatively different antibody responses against SARS-CoV-2 variants. medRxiv., 2021.09.27.21264163 (2022).

11. M. Hoffmann et al., The Omicron variant is highly resistant against antibody-mediated neutralization: Implications for control of the COVID-19 pandemic. Cell. 185, 447–456 (2021).

12. C. O. Barnes et al., SARS-CoV-2 neutralizing antibody structures inform therapeutic strategies. Nature. 588, 682–687 (2020).

13. A. J. Greaney et al., Antibodies elicited by mRNA-1273 vaccination bind more broadly to the receptor binding domain than do those from SARS-CoV-2 infection. Science Translational Medicine. 13(2021).

14. A. J. Greaney et al., Mapping mutations to the SARS-CoV-2 RBD that escape binding by different classes of antibodies. Nat Commun. 12, 1–14 (2021).

15. E. Cameroni et al., Broadly neutralizing antibodies overcome SARS-CoV-2 Omicron antigenic shift. Nature.(2021).

16. L. Liu et al., Striking antibody evasion manifested by the Omicron variant of SARS-CoV-2. Nature.(2021).

17. M. McCallum et al., Structural basis of SARS-CoV-2 Omicron immune evasion and receptor engagement. Science.(2022).

18. Y. Cao et al., Omicron escapes the majority of existing SARS-CoV-2 neutralizing antibodies. Nature.(2021).

19. D. Planas et al., Considerable escape of SARS-CoV-2 Omicron to antibody neutralization. Nature.(2021).

20. A. Aggarwal, et al., SARS-CoV-2 Omicron: evasion of potent humoral responses and resistance to clinical immunotherapeutics relative to viral variants of concern. medRxiv., 2021.12.14.21267772 (2021).

21. L. A. VanBlargan et al., An infectious SARS-CoV-2 B.1.1.529 Omicron virus escapes neutralization by several therapeutic monoclonal antibodies. Nature Medicine.(2022).

22. W. Dejnirattisai et al., Omicron-B.1.1.529 leads to widespread escape from neutralizing antibody responses. Cell. 185, 467–484.e15 (2022).

23. S. Iketani, et al., Antibody Evasion Properties of SARS-CoV-2 Omicron Sublineages. bioRxiv., 2022.02.07.479306 (2022).

24. K. Westendorf, et al., LY-CoV1404 (bebtelovimab) potently neutralizes SARS-CoV-2 variants. bioRxiv., 2021.04.30.442182 (2022).

25. A. Baum et al., Antibody cocktail to SARS-CoV-2 spike protein prevents rapid mutational escape seen with individual antibodies. Science.(2020).

26. C. Hsieh et al., Structure-based design of prefusion-stabilized SARS-CoV-2 spikes. Science. 369, 1501–1505 (2020).

27. T. N. Starr et al., Deep Mutational Scanning of SARS-CoV-2 Receptor Binding Domain Reveals Constraints on Folding and ACE2 Binding. Cell. 182, 1295–1310.e20 (2020).

28. A. J. Schmitz et al., A vaccine-induced public antibody protects against SARS-CoV-2 and emerging variants. Immunity. 54, 2159–2166.e6 (2021).

29. M. A. Tortorici et al., Ultrapotent human antibodies protect against SARS-CoV-2 challenge via multiple mechanisms. Science. 370, 950–957 (2020).

30. T. Li et al., Potent SARS-CoV-2 neutralizing antibodies with protective efficacy against newly emerged mutational variants. Nat Commun. 12, 1–11 (2021).

31. J. Dong et al., Genetic and structural basis for SARS-CoV-2 variant neutralization by a two-antibody cocktail. Nature Microbiology. 6, 1233–1244 (2021).

32. C. Fenwick et al., A highly potent antibody effective against SARS-CoV-2 variants of concern. Cell Reports. 37, 109814 (2021).

33. L. Wang et al., Ultrapotent antibodies against diverse and highly transmissible SARS-CoV-2 variants. Science.(2021).

34. V. Dussupt et al., Low-dose in vivo protection and neutralization across SARS-CoV-2 variants by monoclonal antibody combinations. Nature Immunology. 22, 1503–1514 (2021).

35. Z. Liu et al., Identification of SARS-CoV-2 spike mutations that attenuate monoclonal and serum antibody neutralization. Cell Host & Microbe. 29, 477–488.e4 (2021).

36. D. M. Weinreich et al., REGN-COV2, a Neutralizing Antibody Cocktail, in Outpatients with Covid-19. N. Engl. J. Med. 384, 238–251 (2021).

37. A. Gupta et al., Early Treatment for Covid-19 with SARS-CoV-2 Neutralizing Antibody Sotrovimab. N. Engl. J. Med. 385, 1941–1950 (2021).

38. A. E. Shapiro, R. A. Bender Ignacio, Time to knock monoclonal antibodies off the platform for patients hospitalised with COVID-19. The Lancet Infectious Diseases. 0(2021).

39. C. H. GeurtsvanKessel et al., Divergent SARS CoV-2 Omicron-specific T- and B-cell responses in COVID-19 vaccine recipients. Science Immunology.(2022).

40. C. Wang et al., A conserved immunogenic and vulnerable site on the coronavirus spike protein delineated by cross-reactive monoclonal antibodies. Nat Commun. 12, 1–15 (2021).

41. C. Wang et al., A human monoclonal antibody blocking SARS-CoV-2 infection. Nat Commun. 11, 1–6 (2020).

42. I. Widjaja et al., Towards a solution to MERS: protective human monoclonal antibodies targeting different domains and functions of the MERS-coronavirus spike glycoprotein. Emerg. Microbes Infect. 8, 516–530 (2019).

43. J. Hansen et al., Studies in humanized mice and convalescent humans yield a SARS-CoV-2 antibody cocktail. Science. 369, 1010–1014 (2020).

44. D. Pinto et al., Cross-neutralization of SARS-CoV-2 by a human monoclonal SARS-CoV antibody. Nature. 583, 290–295 (2020).

45. M. Yuan et al., A highly conserved cryptic epitope in the receptor binding domains of SARS-CoV-2 and SARS-CoV. Science. 368, 630–633 (2020).

46. J. Zivanov et al., New tools for automated high-resolution cryo-EM structure determination in RELION-3. eLife. 7, e42166 (2018).

47. A. Punjani, J. L. Rubinstein, D. J. Fleet, M. A. Brubaker, cryoSPARC: algorithms for rapid unsupervised cryo-EM structure determination. Nature Methods. 14, 290–296 (2017).

48. A. Punjani, H. Zhang, D. J. Fleet, Non-uniform refinement: adaptive regularization improves single-particle cryo-EM reconstruction. Nature Methods. 17, 1214–1221 (2020).

49. E. F. Pettersen et al., UCSF Chimera--a visualization system for exploratory research and analysis. J Comput Chem. 25, 1605–1612 (2004).

50. R. Sanchez-Garcia et al., DeepEMhancer: a deep learning solution for cryo-EM volume post-processing. Commun Biol. 4, 1–8 (2021).

51. COSMIC2: A Science Gateway for Cryo-Electron Microscopy Structure Determination, Jul 09-13, 2017(ACM, Proceedings of the Practice and Experience in Advanced Research Computing 2017 on sustainability, success and impact, Jul 09-13, 2017).

52. P. Emsley, K. Cowtan, Coot: model-building tools for molecular graphics. Acta Crystallogr D Biol Crystallogr. 60, 2126–2132 (2004).

53. P. E, et al., SARS-CoV-2 can recruit a heme metabolite to evade antibody immunity. SciAdv. 7, 1–14 (2021).

54. J. Lan et al., Structure of the SARS-CoV-2 spike receptor-binding domain bound to the ACE2 receptor. Nature. 581, 215–220 (2020).

55. L. A. Kelley, S. Mezulis, C. M. Yates, M. N. Wass, M. J. E. Sternberg, The Phyre2 web portal for protein modeling, prediction and analysis. Nature Protocols. 10, 845–858 (2015).

56. P. Emsley, M. Crispin, Structural analysis of glycoproteins: building N-linked glycans with Coot. Acta Crystallogr D Struct Biol. 74, 256–263 (2018).

57. J. J. Headd et al., Use of knowledge-based restraints in phenix.refine to improve macromolecular refinement at low resolution. Acta Crystallogr D Biol Crystallogr. 68, 381–390 (2012).

58. V. B. Chen et al., MolProbity: all-atom structure validation for macromolecular crystallography. Acta Crystallogr D Biol Crystallogr. 66, 12–21 (2010).

59. B. A. Barad et al., EMRinger: side chain-directed model and map validation for 3D cryo-electron microscopy. Nature Methods. 12, 943–946 (2015).

60. J. Agirre et al., Privateer: software for the conformational validation of carbohydrate structures. Nat Struct Mol Biol. 22, 833–834 (2015).

61. J. Agirre, G. Davies, K. Wilson, K. Cowtan, Carbohydrate anomalies in the PDB. Nature Chemical Biology. 11, 303 (2015).

62. E. Krissinel, K. Henrick, Inference of macromolecular assemblies from crystalline state. J Mol Biol. 372, 774–797 (2007).

63. R. A. Laskowski, M. B. Swindells, LigPlot+: Multiple Ligand–Protein Interaction Diagrams for Drug Discovery. J. Chem. Inf. Model. 51, 2778–2786 (2011).

64. T. D. Goddard et al., UCSF ChimeraX: Meeting modern challenges in visualization and analysis. Protein Sci. 27, 14–25 (2018).

65. A. Morin et al., Collaboration gets the most out of software. Elife. 2, e01456 (2013).

66. D. K. Meyerholz, J. C. Sieren, A. P. Beck, H. A. Flaherty, Approaches to Evaluate Lung Inflammation in Translational Research. Vet Pathol. 55, 42–52 (2018).

67. F. Armando, et al., SARS-CoV-2 Omicron variant causes mild pathology in the upper and lower respiratory tract of Syrian golden hamsters (Mesocricetus auratus). (2022).

68. K. Becker et al., Vasculitis and Neutrophil Extracellular Traps in Lungs of Golden Syrian Hamsters With SARS-CoV-2. Frontiers in Immunology. 12, 640842 (2021).

69. B. Bošnjak et al., Intranasal Delivery of MVA Vector Vaccine Induces Effective Pulmonary Immunity Against SARS-CoV-2 in Rodents. Front Immunol. 12, 772240 (2021).

